# Linking molecular pathways and islet cell dysfunction in human type 1 diabetes

**DOI:** 10.1101/2025.02.20.639325

**Authors:** Theodore dos Santos, Xiao Qing Dai, Robert C Jones, Aliya F Spigelman, Hannah M Mummey, Jessica D Ewald, Cara E Ellis, James G Lyon, Nancy Smith, Austin Bautista, Jocelyn E Manning Fox, Norm F. Neff, Angela Detweiler, Michelle Tan, Rafael Arrojo E Drigo, Jianguo Xia, Joan Camunas-Soler, Kyle J Gaulton, Stephen Quake, Patrick E MacDonald

## Abstract

Type 1 diabetes (T1D) is characterized by the autoimmune destruction of most insulin-producing β-cells, along with dysregulated glucagon secretion from pancreatic α-cells. We conducted an integrated analysis that combines electrophysiological and transcriptomic profiling, along with machine learning, of islet cells from T1D donors to investigate the mechanisms underlying their dysfunction. Surviving β-cells exhibit altered electrophysiological properties and transcriptomic signatures indicative of increased antigen presentation, metabolic reprogramming, and impaired protein translation. In α-cells, we observed hyper-responsiveness and increased exocytosis, which are associated with upregulated immune signaling, disrupted transcription factor localization and lysosome homeostasis, as well as dysregulation of mTORC1 complex signaling. Notably, key genetic risk signals for T1D were enriched in transcripts related to α-cell dysfunction, including MHC class I which were closely linked with α-cell dysfunction. Our data provide novel insights into the molecular underpinnings of islet cell dysfunction in T1D, highlighting pathways that may be leveraged to preserve residual β-cell function and modulate α-cell activity. These findings underscore the complex interplay between immune signaling, metabolic stress, and cellular identity in shaping islet cell phenotypes in T1D.

**Highlights:**

- Surviving β-cells in T1D show disrupted electrical function linked to metabolic reprogramming and immune stress.
- Transcripts associated with α-cell dysfunction are enriched in genetic risk alleles for T1D.
- Upregulated MHC class I and impaired nuclear localization of key transcription factors associate with α-cell dysfunction in T1D.
- T1D α-cells exhibit increased hyper-activity, lysosomal imbalance and impaired mTORC1 signaling, which promotes dysregulated glucagon secretion.

## Introduction

Type 1 diabetes (T1D) is defined by a near-complete loss of circulating insulin and dysregulated glucagon secretion. The auto-immune destruction of insulin-producing β-cells in T1D is incomplete, however, as many individuals have detectable levels of circulating insulin or C-peptide^1^ and insulin-positive cells within their pancreas, even after many years of disease^2^. Indeed, circulating C-peptide levels correlate with lower rates of microvascular complications in T1D^3^. Early in T1D, even before clinical onset, β-cells may be dysfunctional^4–7^, and it is debated whether surviving β-cells in clinical T1D display relatively normal^8,9^ or abnormal^10–15^ functional phenotypes. Molecular profiling suggests increased inflammatory and ER stress signatures in β-cells from donors with T1D^10,16–19^.

Circulating glucagon is too high at normoglycemia in T1D, and not appropriately stimulated upon hypoglycemia^20^. Individuals with T1D have elevated glucagon responses to a mixed meal tolerance test^21^ and increased glucagon sensitivity to glucose-dependent insulinotropic polypeptide^22^ (GIP). In vitro, glucagon secretion from islets from donors with T1D is reportedly dysregulated^8^ and appears elevated from islets of autoantibody-positive donors, with increased responsiveness to cAMP-raising stimuli^23^. Molecular profiling reveals increased expression of MHC class I on T1D α-cells^24,25^, consistent with increased inflammatory and immune signaling and elevated ER stress^8,25^, along with potential changes in the expression of α-cell identity markers^17,26^.

Here we investigate the mechanisms of islet cell dysfunction in T1D through linked electrophysiological and transcriptomic analyses. Islets from T1D donors had minimal insulin but retained some β-cells with altered electrophysiological phenotypes, indicating disrupted function. Transcriptomic changes suggested increased antigen presentation (MHC class I), a metabolic shift towards glycolysis, and downregulation of protein translation, indicating metabolic reprogramming linked to autoimmune activity. The α-cells of T1D islets showed hyper-responsiveness and increased activity. Molecular profiling indicated disrupted α-cell identity and regulatory control, and alterations to MHC class I and glucagon compartmentalization, while mTORC1 dysregulation enhanced excitability and exocytosis. Additionally, alterations in lysosomal balance and upregulated proteasomes indicate chronic cellular stress affecting protein turnover. These mechanisms contribute to the hyperactive, glucose-unresponsive phenotype in T1D α-cells. Notably, genetic risk alleles for T1D were enriched in transcripts associated with α-cell dysfunction. Our findings highlight distinct dysfunctional states in T1D α-and β-cells driven by transcriptomic and electrophysiological reprogramming.

## Results

### Islet isolation and cell phenotyping from donors with T1D

Consistent with previous single^9,12,27^ or multi-donor^9,12^ studies, we isolated islets from organ donors with T1D (**Fig 1A**; **Suppl Table 1**). We did not perform density gradient separations but could observe dithizone-positive tissue, albeit at lower levels, suggesting zinc-containing islets with β-cells in T1D (**Fig 1B**). From the T1D donors, α-cell-enriched islets were present in pancreatic tail biopsies and in the isolated islets, where β-cells could be observed sporadically (**Fig 1B**). Insulin content was very low in islets from the T1D donors, while glucagon content was not significantly different from age and sex-matched controls (**Fig 1C**). Secreted insulin was difficult to detect reliably from islets of T1D donors (not shown), while glucagon secretion was hyper-responsive to stimuli (**Fig 1C**) consistent with increased glucagon responses to a mixed meal in T1D^21^ and reports of increased responsiveness of glucagon secretion to cAMP-raising agents from islets of autoantibody-positive donors^23^. Isolated α-cells from T1D donors also showed increased excitability, with higher action potential frequency and amplitude than non-diabetic (ND) controls at 5 mM glucose (**Fig 1D**), and much lower ATP-sensitive K^+^ (K_ATP_) channel current density (**Fig 1E**). Compared to ND α-cells, which showed a significant reduction in total exocytosis from 1 to 5 mM glucose, T1D α-cell exocytosis was not suppressed by elevating glucose (**Fig 1E**).

**Figure 1:**
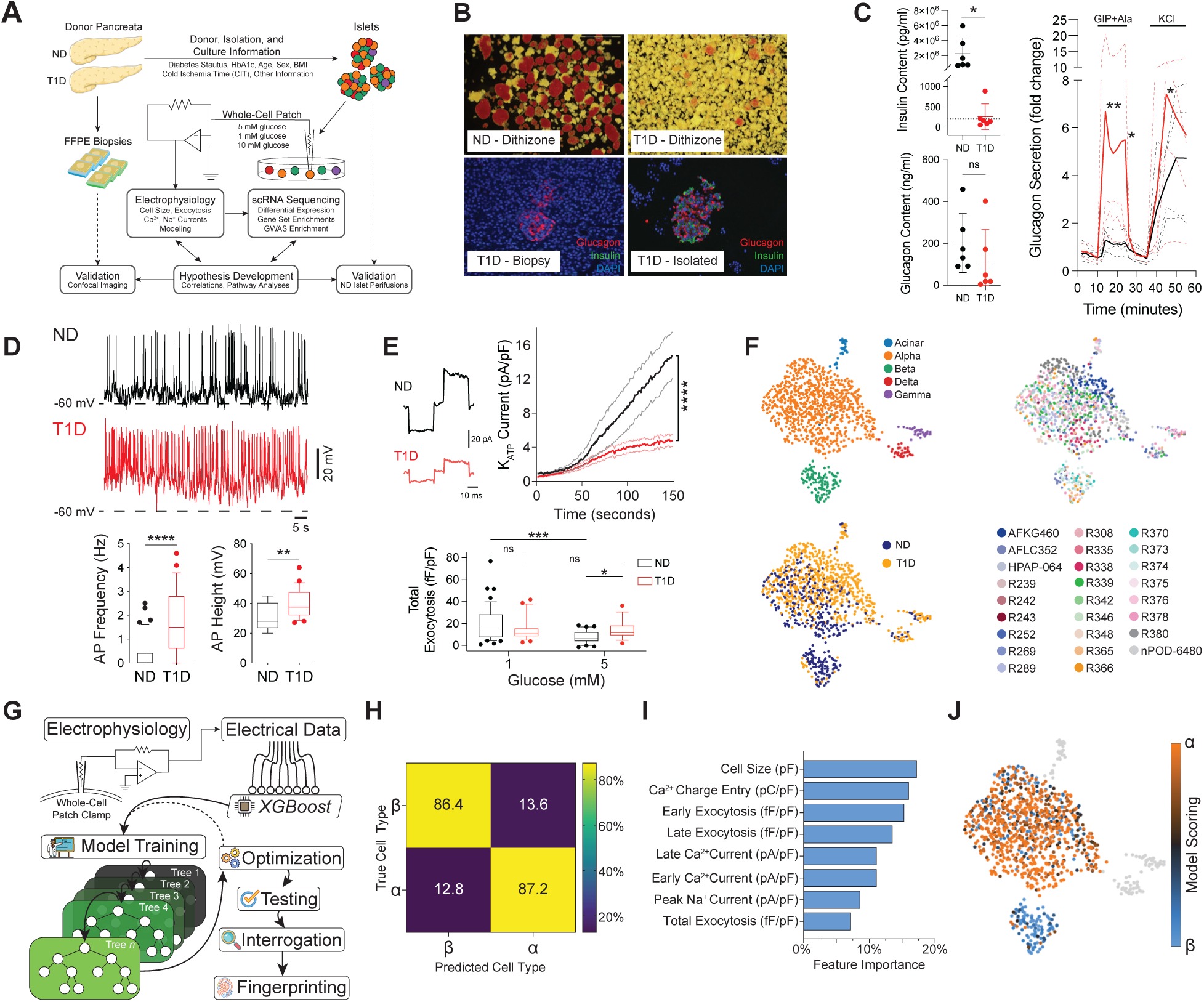
Patch-seq and electrophysiological fingerprinting of human donor islets. (A) Illustration of the isolation and dispersion of human donor islets for patch-seq, followed by the analysis of the single cell transcriptomes and electrophysiology using correlations and modeling approaches, and subsequent validation methods. (B) Dithizone staining of isolated islets from ND and T1D donors, and immunofluorescent staining for glucagon (red), insulin (green), and nuclei (blue) in T1D islets from pancreatic tail biopsy or post isolation. (C) Glucagon and insulin content in islets from ND (n = 4) and T1D (n = 4) donors with glucagon perifusion profiles. (D) Representative actional potential (AP) traces and quantification of their frequency and amplitude (height) in ND (n = 27, 2 donors) and T1D (n = 49, 4 donors) α-cells. (E) Representative K_ATP_ current traces and quantification in ND (n = 42, 4 donors) and T1D (n = 29, 2 donors) α-cells, with normalized total exocytosis at 1mM (ND = 44, 8 donors and T1D = 24, 2 donors) and 5mM glucose (ND = 31, 8 donors and T1D = 18, 2 donors). (F) UMAP of T1D (n=692, 9 donors) and control ND (n=375, 17 donors) patch-seq cells, with overlays of cell type, diabetes status, and donor ID. (G) Illustration of the machine learning classifier model training and testing used to generate scores for electrophysiological fingerprinting. (H) Confusion matrix of model performance at classifying *a priori* data. (I) Feature importance graph showing the extent of dependency on the electrophysiology parameters used by the model for scoring. (J) UMAP from (F) with model scoring overlay of the electrophysiological fingerprinting. ^∗^p < 0.05, ^∗∗^p < 0.01, ^∗∗∗^p < 0.001, and ^∗∗∗∗^p < 0.0001 as indicated using the two-tailed non-parametric Mann-Whitney test (C, D, E), or two-way ANOVA (C, E).

We collected pancreas patch-seq data^17^ from islet cells of donors with and without T1D and identified cell types by integrating these with a larger set of islet cell patch-seq data^17,26^ (**Suppl Table 1**). Leiden clusters with the expression of *GCG*, *INS* and *IAPP*, *SST*, *PPY*, or *PRSS1* and *PRSS2,* grouped together in UMAP space, and were identified as alpha (α), beta (β), delta (δ), gamma (γ), and acinar cells, respectively (**Suppl Fig 1**). Overlay of donor metadata such as age, BMI, HbA1c (%), sex, diabetes status, and diabetes duration suggested good integration within a cell type cluster (**Suppl Fig 2**). Because organ donors with T1D are typically younger than the standard donor pool, subsequent analyses used a cohort of controls matched for age, sex, body mass index (BMI), and cold ischemic time (CIT) (**Suppl Fig 3**). The resulting 1067 cells from 17 ND and 9 T1D donors retained clustering by cell type (**Fig 1F, Suppl Fig 4**) irrespective of diabetes status (**Fig 1F**), and showed good integration (**Sup Fig 5**).

To provide a metric representing overall canonical electrical behavior, we performed electrophysiological fingerprinting^26,28^. We created an advanced machine learning model (**Fig 1G**) trained on a larger data set (α-cells: 258, β-cells: 290) of the matched ND electrical data expanded with patched cells whose identities were confirmed by immunostaining for insulin or glucagon from www.humanislets.com^29^ (**Sup Table 1**). The model performs with an accuracy of ∼85% in correctly identifying α- and β-cells on electrophysiologic data alone, which is not due to trivial majority class prediction (**Fig 1H**), with a balanced reliance on multiple electrical parameters (**Fig 1I**) and was not impacted by donor characteristics (**Sup Fig 6**). This allows the assignment of probabilities to α-cell (α-score) or β-cell (β-score) identities, which are indicative of their overall electrical phenotypes and can subsequently be overlaid on top of the transcriptomic phenotypes (**Fig 1J**).

### Transcriptomic alterations and loss of normal electrical function in T1D β- and α-cells

While the number of T1D β-cells is relatively small (n=16, 4 donors), they possessed notably different transcriptomic and electrical profiles compared to ND (n=104, 14 donors) at 5 mM glucose. Differential expression analysis (DEA) and subsequent Gene Set Enrichment Analysis (GSEA) (**Suppl Table 2**) revealed markers (**Fig 2A**) and pathways (**Fig 2B**) that included some expected findings such as up-regulation of antigen presentation related to MHC class I^30,31^ and interferon-gamma^32,33^, and down-regulation of protein translation^31^. Notably, T1D β-cells show enrichment of markers (**Fig 2A**) and pathways (**Fig 2B**) involved in glycolysis^31^ (**Fig 2A, *purple***), and downregulation of markers involved in pyruvate-related metabolism (**Fig 2A, *blue***) and mitochondrial respiration (**Fig 2B**), suggesting a metabolic shift towards non-oxidative glucose metabolism. Except for *INS*, transcripts of T1D autoantibody targets such as *IAPP, GAD2, IGF2, and APP* were upregulated. Functionally, T1D β-cells show increased exocytosis and larger inward (i.e. more negative) voltage-activated Ca^2+^ currents compared to controls (**Fig 2C**). Insulin secretion from β-cells of T1D donors has been reported to be either normal^8,9^ or impaired^12,27^ although function could perhaps be restored after several days in culture^13^. Notably, the scoring (β-score) by our classifier model was significantly decreased in T1D β-cells, indicative of substantially altered β-cell electrical function (**Fig 2C**).

**Figure 2:**
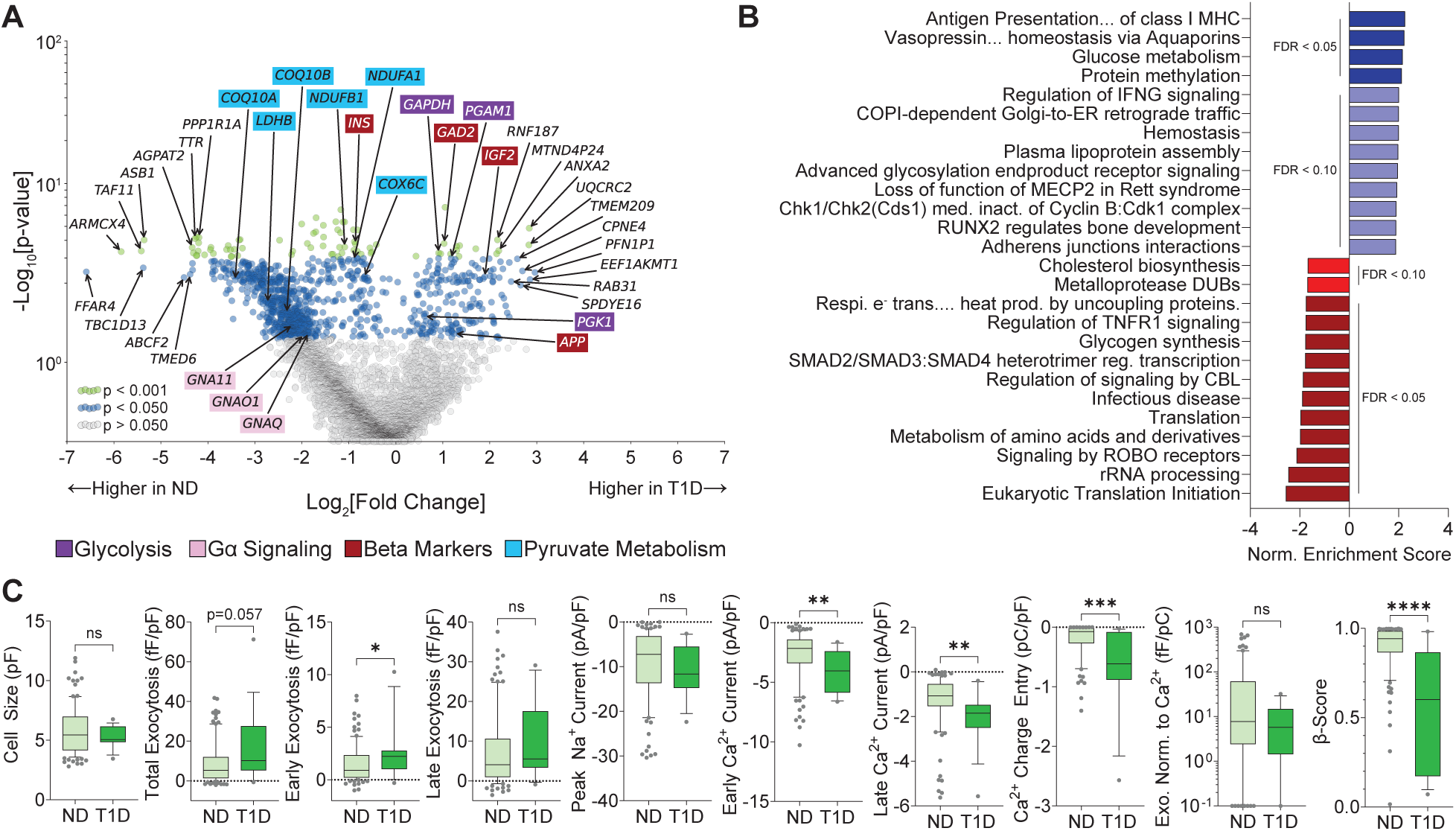
Patch-seq of T1D β-cells at 5mM glucose. (A) Differential expression analysis of the patch-seq β-cells, represented as a volcano plot showing transcripts with increased (right) and decreased (left) expression in T1D (n=16, 4 donors) compared to ND (n = 104, 14 donors). Uncolored annotations show the top 10 and bottom 10 most differentially expressed transcripts. Colored annotations highlight notable transcripts involved with glycolysis (purple), pyruvate metabolism (turquoise), Gα signaling (pink), and auto-antigen beta markers that elicit autoantibody responses in T1D (red). (B) Gene set enrichment analysis using the scores from the differential expression analysis in (A) as weighting, referenced against the Reactome pathway database. Pathways upregulated in T1D are shown in blues, while those downregulated are shown in reds. (C) Comparison of the electrophysiology and model scoring obtained from patch-seq β-cells during whole-cell patch-clamp at 5mM glucose. ND (14 donors) and T1D (4 donors) n-values for cell size = 104, 16; total exocytosis = 102, 16; early exocytosis = 102, 15; late exocytosis = 102, 15; peak sodium current = 104, 16; early Ca^2+^ current = 101, 16; late Ca^2+^ current 101, 16; Ca^2+^ charge entry = 83, 16; exocytosis normalized to Ca^2+^ = 80, 16; model scoring = 104, 16. ^∗^p < 0.05, ^∗∗^p < 0.01, ^∗∗∗^p < 0.001, and ^∗∗∗∗^p < 0.0001 as indicated using the two-tailed non-parametric Mann-Whitney test. Outliers were determined based on |z-score| >3 and excluded from comparison and statistical analysis.

Like the β-cells, T1D α-cells (n = 596, 9 donors) are enriched (**Suppl Table 2)** in antigen presentation markers (**Fig 3A, blue**) and pathways (**Fig 3B**) related to MHC class I compared to matched controls (n = 248, 17 donors). We also observed an increase in pathways related to incretin secretion^34,35^. Our previous work in type 2 diabetes (T2D) showed that α-cell dysfunction is linked to lineage and maturity state^26^, and we observe an enrichment of these markers in T1D α-cells (**Fig 3A, *orange***), corroborating with previous findings^25^. Key receptors of endocrine and paracrine signaling that modulate glucagon secretion are also upregulated (**Fig 3A, *turquoise***), including the GIP receptor (*GIPR*), with an enrichment in Gα (s) signaling and signaling by GPCR (**Fig 3B**). Additionally, pathways for gluconeogenesis, carbohydrate metabolism, TCA cycle, and lipid metabolism are enriched in T1D α-cells (**Fig 3B**). Downregulated pathways of note included multiple cytokine-related pathways and mammalian/mechanistic target of rapamycin complex 1 (mTORC1) (**Fig 3B**). We detected an enrichment of transcripts differentially expressed in T1D α-cells among genes with T1D genetic association, as well as genes directly regulated by T1D associated variants from islet expression QTL (eQTL) data (**Fig 3C**).

**Figure 3:**
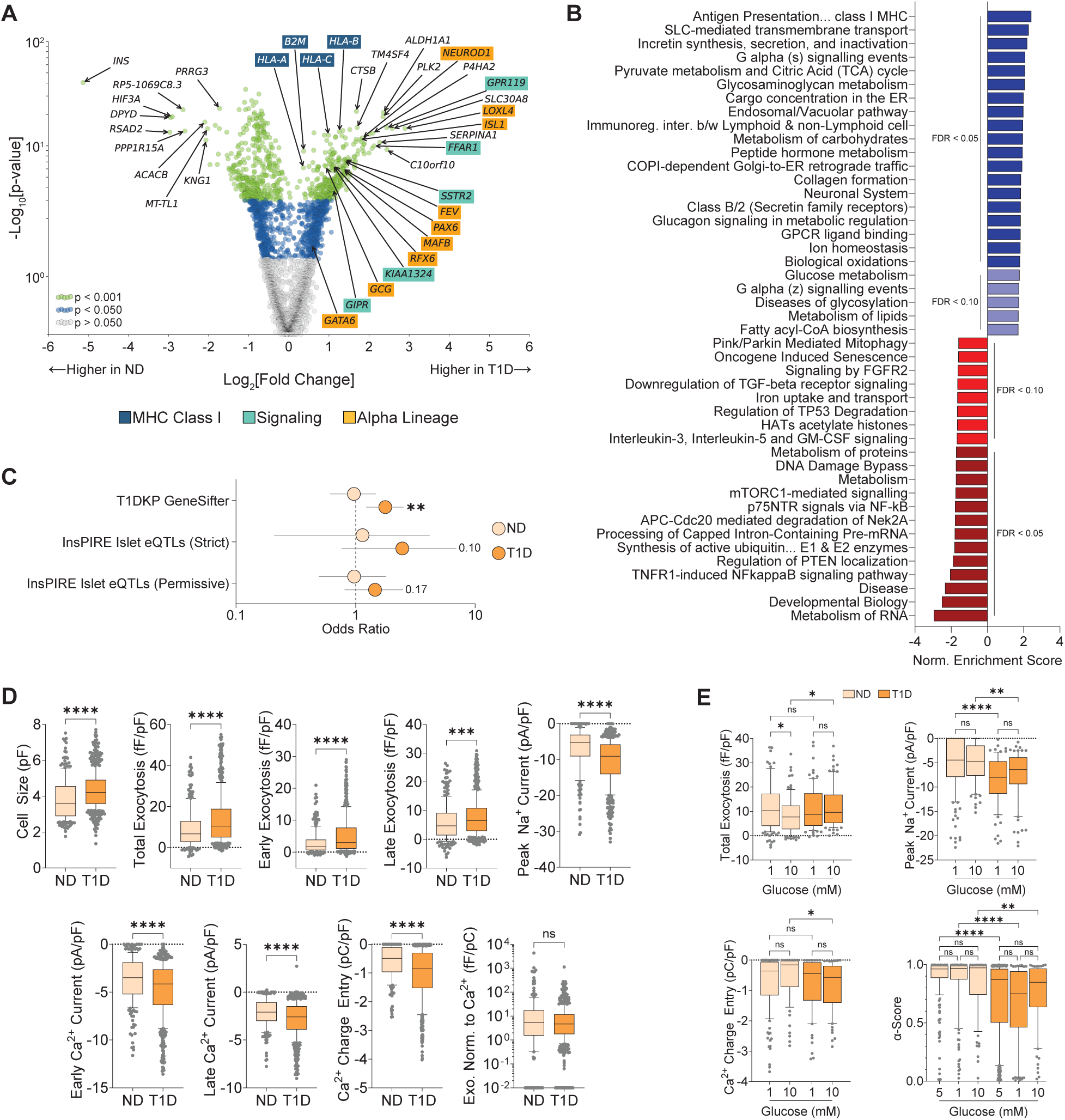
Patch-seq of T1D α-cells. (A) Differential expression analysis of the patch-seq α-cells, represented as a volcano plot showing transcripts with increased (right) and decreased (left) expression in T1D (n=596, 9 donors) compared to ND (n = 248, 17 donors). Uncolored annotations show the top 10 and bottom 10 most differentially expressed transcripts. Colored annotations highlight notable transcripts involved with MHC class I antigen presentation (blue), signaling (turquoise), and α-cell markers of lineage and immaturity state (orange). (B) Gene set enrichment analysis using the scores from the differential expression analysis in (A) as weighting, referenced against the Reactome pathway database. Pathways upregulated in T1D are shown in blues, while those downregulated are shown in reds. (C) Odds ratio for the enrichment of T1D risk genes in differentially expressed transcripts in T1D α-cells. Risk genes were identified from the T1D Knowledge Portal and islet eQTLs for T1D-associated variants. Error bars represent the 95% confidence interval of the odds ratio. (D) Comparison of the electrophysiology obtained from the patch-seq α-cells during whole-cell patch-clamp at 5mM glucose. ND (17 donors) and T1D (9 donors) n-values for cell size = 245, 585; total exocytosis = 243, 588; early exocytosis = 241, 583; late exocytosis = 246, 584; peak sodium current = 238, 577; early Ca^2+^ current = 236, 580; late Ca^2+^ current 236, 577; Ca^2+^ charge entry = 205, 531; exocytosis normalized to Ca^2+^ = 195, 529. (E) Comparison of the electrophysiology and model scoring at differing glucose concentrations obtained from patch-seq α-cells during whole-cell patch-clamp. At 1mM glucose, ND (13 donors) and T1D (4 donors) n-values for cell size = 156, 81; total exocytosis = 154, 81; peak sodium current = 154, 78; Ca^2+^ charge entry = 149, 75. At 10mM glucose, ND (10 donors) and T1D (4 donors) n-values for cell size = 85, 80; total exocytosis = 84, 80; peak sodium current = 84, 81; Ca^2+^ charge entry = 77, 70. Model scoring of electrophysiology recorded at 5mM, 1mM, and 10mM from patch-seq α-cells of ND matched controls (17, 13, 10 donors respectively) and T1D (9, 4, 4 donors respectively). ND and T1D n-values for model score at 5mM = 248, 596; 1mM = 158, 82; 10mM = 85, 81. ^∗^p < 0.05, ^∗∗^p < 0.01, ^∗∗∗^p < 0.001, and ^∗∗∗∗^p < 0.0001 as indicated using Fisher’s Exact Test (C), two-tailed non-parametric Mann-Whitney test in (D) and using the non-parametric Kruskal-Wallis test with Dunn’s correction (E). Outliers were determined based on |z-score| >3 and excluded from comparison and statistical analysis (D and E).

The T1D α-cells are significantly larger than ND α-cells and show greater exocytosis with increased inward Ca^2+^ and Na^+^ currents at 5 mM glucose, even when normalized to cell size (**Fig 3D**). This suggests broad alterations in electrical phenotypes via enhanced excitability and exocytosis, as initially noted above (**Fig 1D-E**), and is corroborated by a decrease in α-score from our classifier, suggesting a loss of canonical electrical behavior (**Fig 3E**). Although α-cell glucagon release is regulated by paracrine signals, α-cell secretion can be impacted by glucose directly^26,36^. We confirmed the reduction in exocytosis and inward Ca^2+^ currents in ND α-cells at high glucose (10mM); however, T1D α-cells fail to show these glucose-mediated changes, while also possessing elevated inward Na^+^ currents (**Fig 3E, Suppl Fig 7**). Scoring from our classifier model is largely glucose-independent, with T1D α-cells showing a significant decrease in α-scores irrespective of glucose levels, suggesting that the loss of electrical phenotype is consistent across glucose concentrations (**Fig 3E, Suppl Fig 7**).

### Diverse markers and pathways linked to elevated exocytosis in T1D α-cells

To identify pathways in T1D α-cells closely linked with their functional deficits, we correlated transcript expression with normalized total exocytosis (here on referred to as ‘exocytosis’; **Suppl Table 3**). Amongst the top 100 significant correlates (**Fig 4A**) were transcripts associated with islet health and diabetes. These include *GC* (vitamin D binding protein), known to contribute to glucagon secretion via actin remodeling and metabolic stress adaptation^37–39^, and *LOXL4* (lysyl oxidase like 4) which is linked to heterogeneity in secretory behavior^17^ and recently discovered to contribute to immune-evasion by suppressing the activity of macrophages and CD8+ T-cells^40,41^. *IFI6* (interferon-alpha inducible protein 6), an ER-residing interferon effector that suppresses flavivirus replication^42^ and previously found to be highly upregulated in T1D^43^, and the suppressor of mitochondrial mediated apoptosis *APIP*^44^ were also positively correlated with exocytosis. Additional correlates include *KCNK16* (TALK-1), a two-pore-domain K^+^ channel whose genetic variants have been closely associated with β-cell dysfunction^45^ and recently linked to increased glucagon secretion in MODY models^46^, and *GIPR* which enhances glucagon secretion from α-cells^22,47^ and may contribute to the increased glucagon response in **Fig 1C**. *MPC1* (Mitochondrial Pyruvate Carrier 1), a mitochondrial carrier that transports pyruvate, an inducer of glucagon secretion^48^, across the mitochondrial membrane into the TCA cycle, also positively correlated with exocytosis.

**Figure 4:**
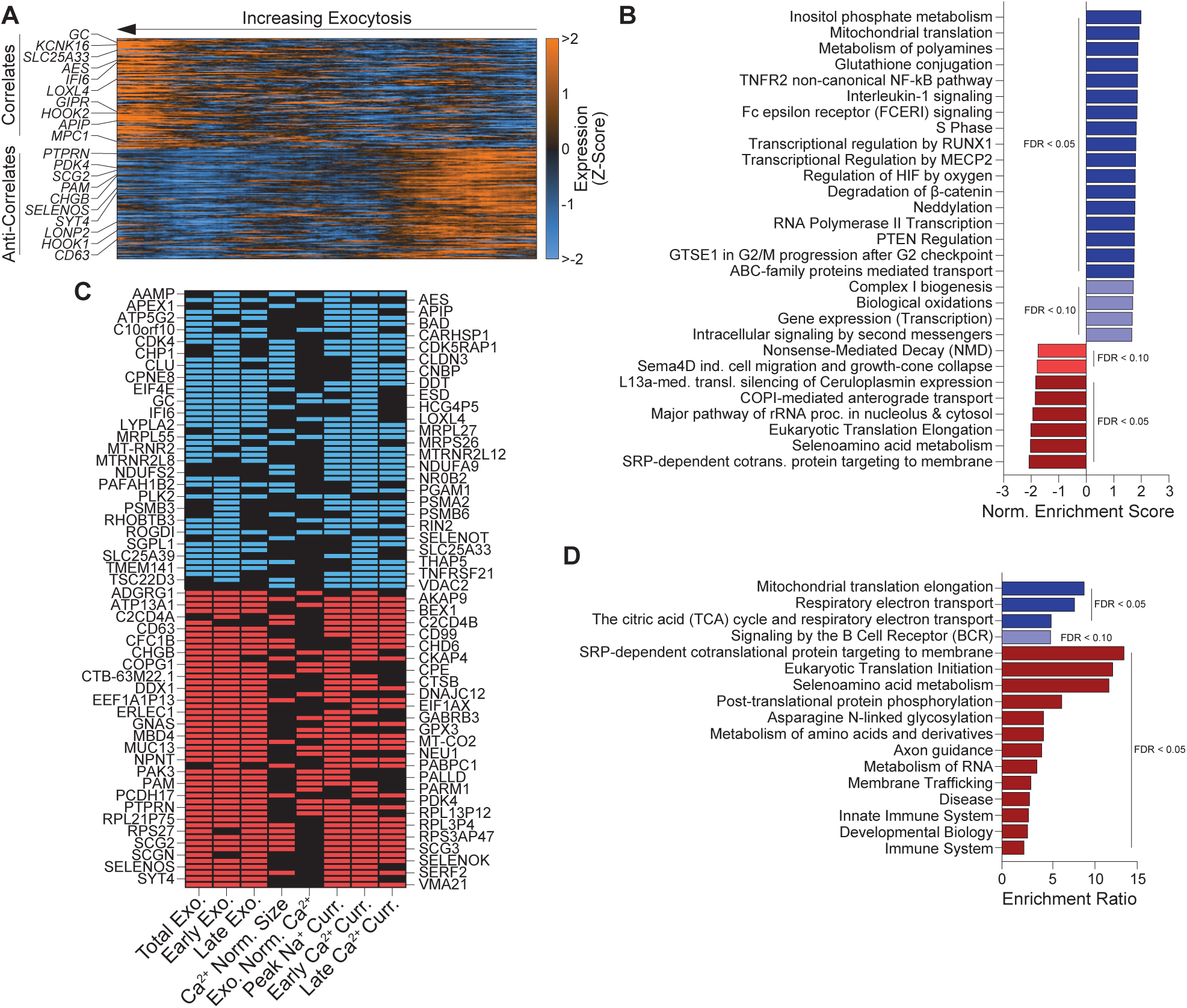
Linking transcript expression with electrophysiology in T1D patch-seq α-cells at 5mM glucose. (A) Heat map representing the top transcripts that correlate and anti-correlate with normalized total exocytosis, if expressed, in at least 50% of T1D α-cells at 5mM glucose. Expression is represented as a Z-score with smoothing (n=50), and cells are sorted by increasing exocytosis from right to left. (B) Gene set enrichment analysis using the expression vs normalized total exocytosis correlation coefficients as weighting, referenced against the Reactome pathway database. Pathways correlating with exocytosis in T1D α-cells are shown in blues, while those anti-correlating are shown in reds. (C) Top transcripts that significantly (p<0.05) correlate (blue) or anti-correlate (red) with secretory electrical behavior across 3 or more recorded electrophysiological parameters. (D) Over representation analysis of transcripts that significantly (p<0.05) correlate (blues) or anti-correlate (red) with secretory electrical behaviour across 3 or more recorded electrophysiological parameters, referenced against the Reactome pathway database.

Anti-correlates include *PDK4* (Pyruvate Dehydrogenase Kinase 4), which reduces pyruvate entry into the TCA cycle by inhibiting pyruvate dehydrogenase in α-cells ^48^ and regulates glucagon secretion^49^, and *HOOK1* (Hook Microtubule Tethering Protein 1) which facilitates Rab5-tagged early endosome transport^50^ and may be linked to endo-lysosome formation that mediates glucagon degradation, lowering secretory capability^51^. Other anti-correlates include *CD63* (Tetraspanin-30) which lowers secretion in β-cells via lysosomal mediated degradation of insulin granules^52^, and *SYT4* (Synaptotagmin 4), which reduces insulin secretion by competing with Ca^2+^ sensitive Synaptotagmin 7^53^. We also detected markers that abrogate cellular stress, such as *SELENOS* (Selenoprotein S), a reductase^54^ that mitigates oxidative and ER related stress^55^, and *LONP2* (Lon Peptidase 2) which degrades oxidized proteins at the peroxisome, alleviating oxidative stress and cellular aging^56,57^. Other negatively correlates of T1D α-cell exocytosis were the granins, *CHGB* (Chromogranin B) and *SCG2* (Secretogranin II), which are well studied in islet cells^58^, and transcripts primarily characterized in β-cells such as *PAM* (Peptidylglycine Alpha-Amidating Monooxygenase), which regulates insulin content and secretion in diabetes^59,60^, as well as *PTPRN* (Protein Tyrosine Phosphatase Receptor Type N) - the T1D-autoantigen IA-2^23^.

We next identified pathways that correlate with T1D α-cell exocytosis (**Fig 4B, Suppl Table 3**), some of which suggest that the immune environment of the T1D islet is influencing α-cell electrical behavior. Interleukin-1, a chronic marker of T1D that also determines the extent of auto-inflammation^61^, and its signaling pathway correlates with inappropriately elevated α-cell exocytosis in T1D. We also detected the TNFR2 non-canonical NF-kB pathway, which is considered a slow and persistent activation pathway for NF-kB, paralleling the chronic nature of T1D^62^. Notably, transcripts contributing to the enrichment of these two pathways in our analysis were also enriched in proteasomal components, which are upregulated in inflammatory conditions^63^. Next, we identified transcripts that correlated with multiple electrophysiological parameters (**Fig 4C, Suppl Table 4**), recapturing many of the markers trending with total exocytosis from **Fig 4A**. Subsequent Over Representation Analysis (ORA) identified pathways enriched for transcripts with broad correlations across electrophysiology parameters (**Fig 4D, Suppl Table 4**), suggesting mitochondrial respiration as a possible contributor to the altered electrical behavior, which we previously associated with α-cell dysfunction in T2D^26^.

### Linking electrophysiological phenotypes to molecular pathways and genetic risk in T1D α-cells

While exocytosis measurements are representative of vesicle-mediated secretion, the α-scores provide an integrated metric for the overall electrical behavior of α-cells (**Fig 1G-J**). Amongst the top correlates of α-scores (**Fig 5A, Suppl Table 4**) were multiple transcripts involved in the proteasome (*PSMA6*, *PSMB6, PSMF1, PSMD12, PSMD7,* and *PSMC3)*, suggesting that T1D α-cells with more canonical electrical identity may have higher proteasomal activity, which could help alleviate ER-stress and unfolded protein responses induced by the auto-immune nature of T1D^25^. Notably, other markers involved mTORC1 signaling such as *LAMTOR1* and *LAMTOR2* (Late Endosomal/Lysosomal Adaptor, MAPK and mTOR Activator 1, 2) which are required for the assembly of the mTORC1 activation complex on lysosomes^64^, and *CCT4* (Chaperonin Containing TCP1 Subunit 4), a chaperone that stabilizes the mTORC1-induced disassembly of the lysosomal V-ATPase, suppressing lysosomal acidification and autophagy^65^. *SUMO1* (Small Ubiquitin Like Modifier 1) also positively correlates with α-score and has previously been shown to influence cAMP-mediated exocytosis in α-cells^66^, as well as suppressing NF-κB mediated immune signaling by increasing the half-life of its inhibitor, IκBα (*NFKBIA)*^67,68^.

**Figure 5:**
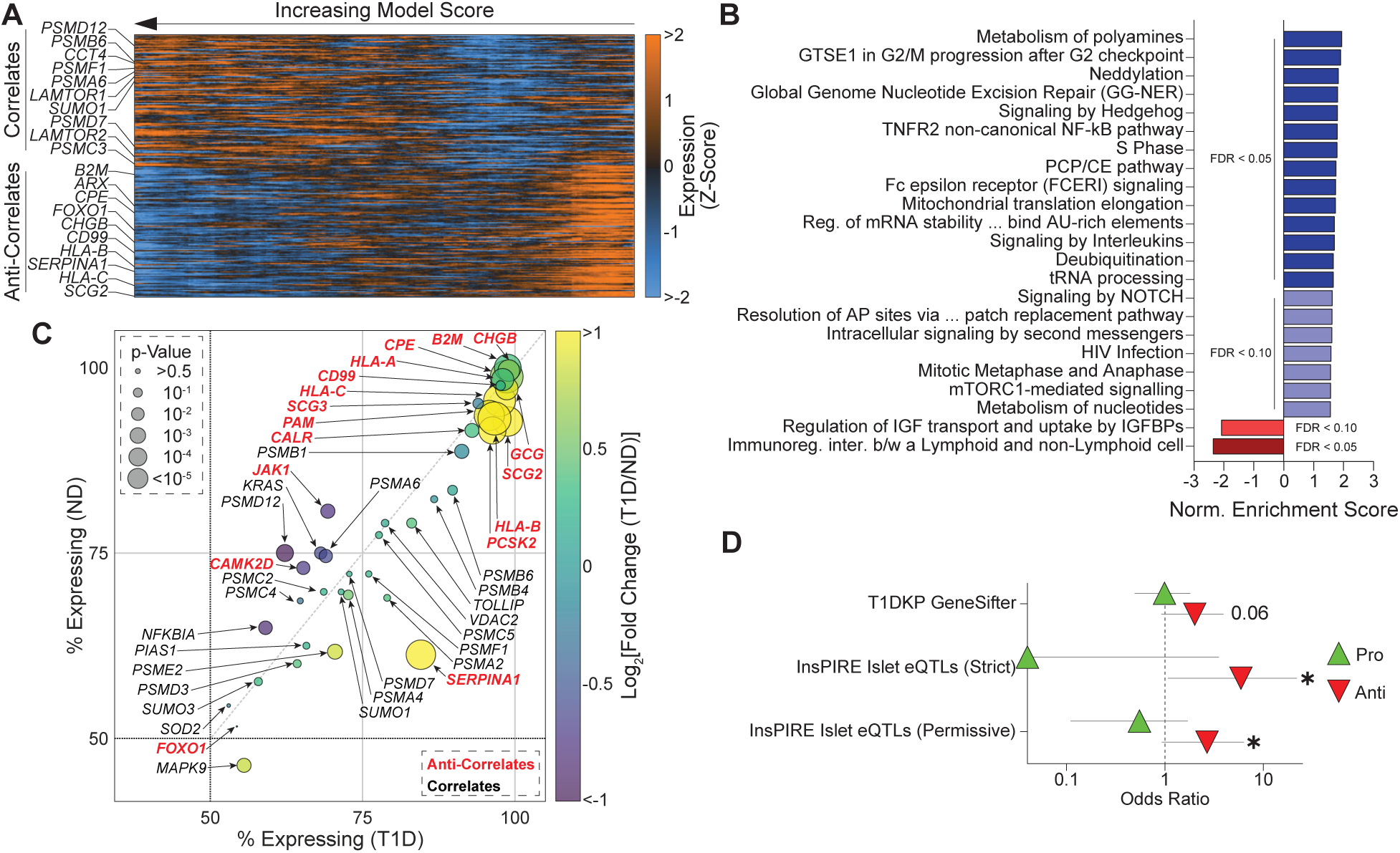
Linking transcript expression with model scoring in T1D patch-seq α-cells at 5mM glucose. (A) Heat map representing the top transcripts that correlate and anti-correlate with α-score, if expressed, in at least 50% of T1D α-cells at 5mM glucose. Expression is represented as a Z-score with smoothing (n=50), and cells are sorted by increasing model score from right to left. (B) Gene set enrichment analysis using the expression vs α-score correlation coefficients as weighting, referenced against the Reactome pathway database. Pathways correlating with α-score in T1D α-cells are shown in blues, while those anti-correlating are shown in reds. (C) Bubble plot of differential expression and proportion of abundance in ND and T1D of notable transcripts that correlate (black) or anti-correlate (red) with α-score in T1D α-cells. (D) Odds ratio for the enrichment of T1D risk genes in transcript expression vs model score correlates (Pro, green triangles) and anti-correlates (Anti, red triangles) in T1D α-cells. Risk genes were identified from the T1D Knowledge Portal and islet eQTLs for T1D-associated variants. Error bars represent the 95% confidence interval of the odds ratio. ^∗^p < 0.01 as indicated using Fisher’s Exact Test.

Top transcripts that negatively correlate with α-score include MHC-1 antigen presentation (*B2M, HLA-B, HLA-C)*, the α-cell transcription factor *ARX* (Aristaless Related Homeobox) which was previously associated with α-cell dysfunction in T2D^26^, and its associated enhancer *FOXO1*^69^ (Forkhead Box O1). *SERPINA1* (Serpin Family A Member 1), a serine protease inhibitor, has been linked to immune and cellular stress responses^70,71^, but has not been directly studied in α-cells under autoimmune stress. Other negative correlates include the granins *SCG2* and *CHGB*^58^, and pro-glucagon processing enzyme *CPE*^51^ (Carboxypeptidase E), whose knockdown leads to increased basal secretion and loss of stimulated secretion in an α-cell culture model^72^.

GSEA of the α-score correlations (**Fig 5B, Suppl Table 4**) revealed several pathways primarily enriched by proteasomal transcripts, including interleukin signaling and TNFR2 non-canonical NF-kB pathways, which we also detected as trending with increased exocytosis (**Fig 4B**). We also detected pathways related to mTORC1-mediated signaling, whose components were previously detected as some of the top trending correlations (**Fig 5A**). Notable positive correlates also included immune signaling effector suppressors *PIAS1*^73^ (Protein Inhibitor of Activated STAT 1), *NFKBIA*^68^, and *TOLLIP* (Toll Interacting Protein), a negative regulator of NF-κB and linked to endo-lysosomes^74^ (**Suppl Table 4**). Though few, pathways that anti-correlate with α-score (i.e. trend with worsening electrical identity) involved insulin growth factor pathways that were enriched for the granins (*SCG2*, *SCG3*, *CHGB*) and *SERPINA1,* and immune signaling, consisting of antigen presentation components (*HLAs, B2M*) and *CD81* (TSPAN28), a recently discovered marker for β-cells displaying dedifferentiation, immaturity, and stress^75^. As many of the notable markers detected in **Fig 5A** and **5B** could be linked to α-cell electrical identity in T1D, we next plotted their differential expression as a dot plot to understand the relevance of their trend with α-score in the context of changes in expression in ND vs T1D (**Fig 5C**). We find that most of the transcripts that negatively correlate with α-score also have significantly higher expression in T1D (**Fig 5C, *red***), and *vice versa* for the positive correlates (**Fig 5C, *black***). Last, we further validated the relevance of these correlative hits by cross-referencing them against genes with evidence for T1D genetic association. We discovered that the negative correlates of α-score significantly overlapped genes directly regulated by T1D risk variants in islet eQTL data (**Fig 5D**), linking together genetic predisposition, variation in transcriptomic expression, and electrical dysfunction.

### Increased inflammatory and immune responses and reduced nuclear import of α-cell identity factors in T1D

We grouped major hits in our correlation results based on their physiological roles to further identify possible mechanisms contributing to α-cell dysfunction in T1D (**Sup Fig 8, Suppl Table 5**). The combined DEA and correlative analysis of MHC-related transcripts is presented in **Fig 6A**. NF-kB is a major driver for the constitutive and induced expression of MHC class I^76^, and we observe that its inhibitor *NFKBIA* (IκBα) is positively associated with α-score and significantly downregulated in T1D. Also, MHC class I transcripts significantly upregulated in T1D consistently trend with worsening α-scores. Glucagon processing and its secretion are highly dependent on secretory granule dynamics, intracellular trafficking, and sorting of vesicle compartments in the α-cells^51,77^, processes shared by MHC class I antigen processing and presentation trafficking to the cell surface^78,79^. Previous work shows an enriched expression of MHC class I on α-cells^24^ whose expression is upregulated further in T1D^25^ and in aging-related cellular stress^80^. Thus, MHC class I upregulation may impact glucagon secretion by altering its sorting and localization in T1D α-cells. 3D confocal microscopy on pancreas sections shows a clear increase in glucagon co-localization with MHC class I signal in T1D donors compared with matched controls (**Fig 6B-C**).

**Figure 6:**
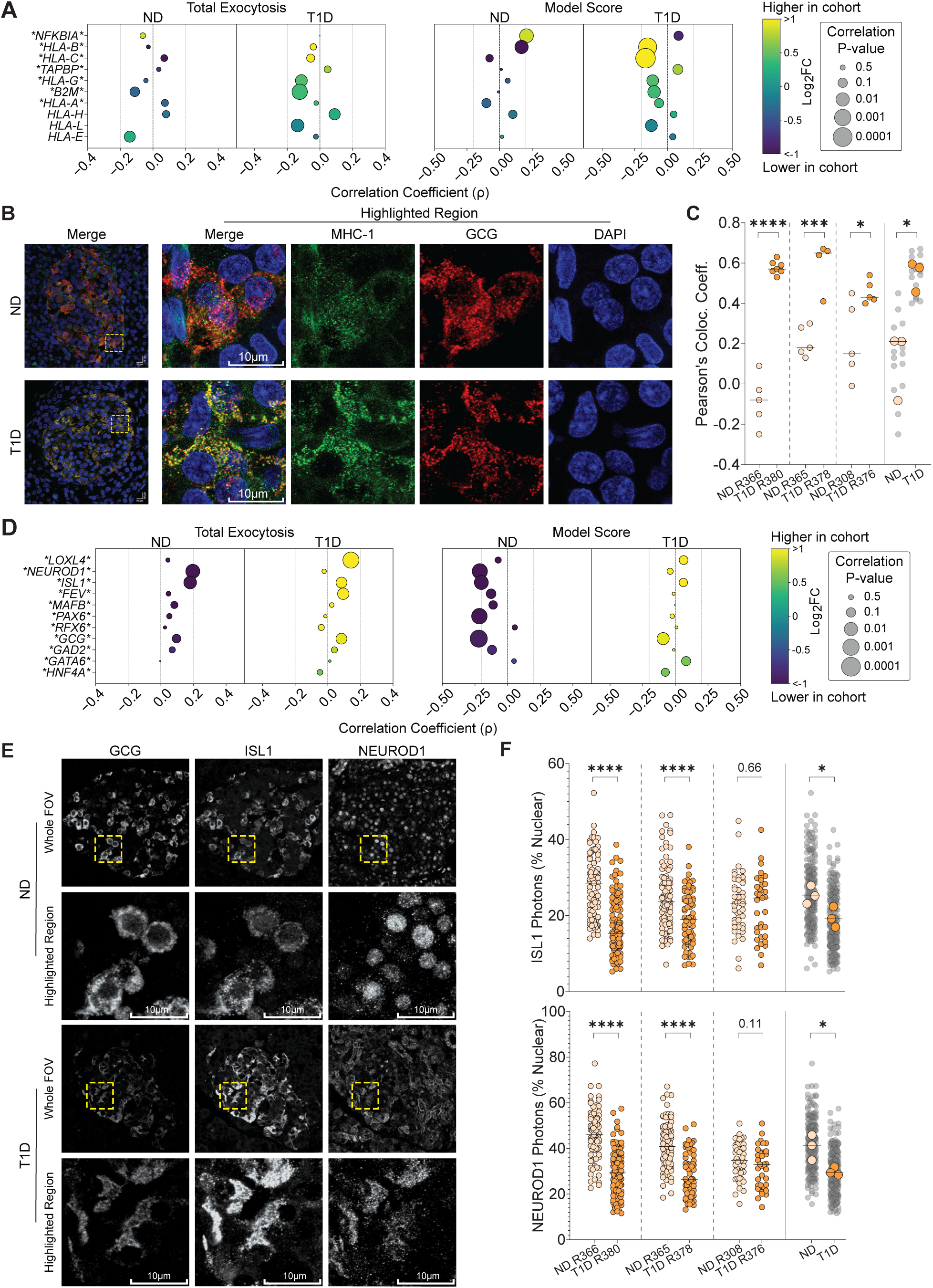
Localization of MHC-1 and α-cell transcription factors ISL1 and NEUROD1 in T1D and matched ND donor biopsies. (A) Bubble plot of notable transcripts involved with antigen presentation, their differential expression in T1D vs ND, and their correlative relationship with total exocytosis and α-score in α-cells at 5mM glucose. Transcript names flanked by asterisks indicate significant (p<0.05) differential expression. (B) Representative images of MHC-1 staining *in situ* from pancreas biopsies of patch-seq donors, attained by immunofluorescence confocal microscopy, presented in pseudo-colors for visually assessing colocalization (yellow). (C) Quantification of MHC-1 and glucagon signal colocalization. Each data point represents a single field of view of an islet from ND (n = 15, 3 donors) and T1D (n = 16, 3 donors) biopsies. ^∗^p < 0.05, ^∗∗∗^p < 0.001, and ^∗∗∗∗^p < 0.0001 as indicated, using the unpaired t-test. Horizontal line markers indicate median. (D) Bubble plot of notable transcripts involved with α-cell lineage and maturity, their differential expression in T1D vs ND, and their correlative relationship with total exocytosis and α-score in α-cells at 5mM glucose. Transcript names flanked by asterisks indicate significant (p<0.05) differential expression. (E) Representative images of ISL1 and NEUROD1 staining *in situ* from pancreas biopsies of patch-seq donors, attained by immunofluorescence confocal microscopy, and presented in greyscale for better contrast. (F) Quantification of ISL1 and NEUROD1 signals that were nuclear in glucagon positive cells. Each data point represents a single cell obtained from ND (n=231, 3 donors) and T1D (n=231, 3 donors) biopsies. ^∗^p < 0.05 and ^∗∗∗∗^p < 0.0001 as indicated, using the unpaired t-test. Horizontal line markers indicate median.

The inflammation and innate immune signaling indicated by these results can lead to changes in homeostatic nuclear permeability due to immunity-related signal transducers, such as NF-κB and STATs, altering nuclear pore components and monopolizing import machinery^81^. This may impair nuclear access for key transcription factors that dictate α-cell identity. For instance, the upregulation of α-cell lineage markers that drive maturation, such as transcription factors *ISL1* and *NEUROD1*, have been previously reported in T1D^25^, and in T2D were associated with dysfunctional α-cell secretory behavior^26^. In our data, these transcripts trend with increasing exocytosis in both the T1D and ND cohort, but in the latter, show a consistent trend with lower α-scores (**Fig 6D**). Indeed, such lineage markers are upregulated in immature α-cells to drive (re-)maturation. However, this trend is lost in the T1D α-cells that show gross electrical dysfunction, despite their transcriptomic upregulation. Thus, we questioned whether the effects of ISL1 and NEUROD1 upregulation were somehow attenuated in T1D α-cells. Using biopsies from the same donors as in Fig 6A-C, we stained for ISL1 and NEUROD1 (**Fig 6E**) and quantified the percentage of cellular signal with nuclear localization in the glucagon-positive cells (**Fig 6F**). Although we observed heterogeneity in nuclear localization within and between donors, nuclear localization was much higher in ND compared to T1D (**Fig 6E, 6F**). Further, irrespective of diabetes status, there was a significant linear trend between the nuclear signals for ISL1 and NEUROD1 (**Suppl Fig 9**). The findings suggest that despite the observed upregulation of α-cell lineage and immaturity markers *NEUROD1* and *ISL1*, they may fail to drive or restore α-cell maturity in T1D due to a general attenuation in nuclear localization.

### T1D α-cell lysosomal disorder with links to reduced mTORC1

The mTORC1 complex is involved in a myriad of cellular pathways^82,83^, with crucial roles in regulating lysosome activity and autophagy^64,65,84^. Key components of the mTORC1 complex trend with lower exocytosis in ND, which is lost in T1D, but still show a positive trend with α-score irrespective of diabetes status, suggesting its importance for canonical behavior (**Fig 7A**). However, these components are significantly downregulated in T1D α-cells (**Fig 7A**). Protein-protein interaction network analysis revealed that amino acid metabolism and regulatory components of mTORC1 activity were significantly abrogated as well (**Fig 7B**). To mimic the consequences of downregulating mTORC1 in T1D α-cells and explore its potential impact on glucagon secretion, we treated ND islets with the potent and highly specific mTOR inhibitor, Torin-2, and performed dynamic secretion assays at 5 mM glucose (**Fig 7C**). We observed that the secretion of glucagon and to a smaller degree, insulin (not shown), were increased upon mTOR inhibition, suggesting that reduced mTORC1 signaling leads to hypersecretion.

**Figure 7:**
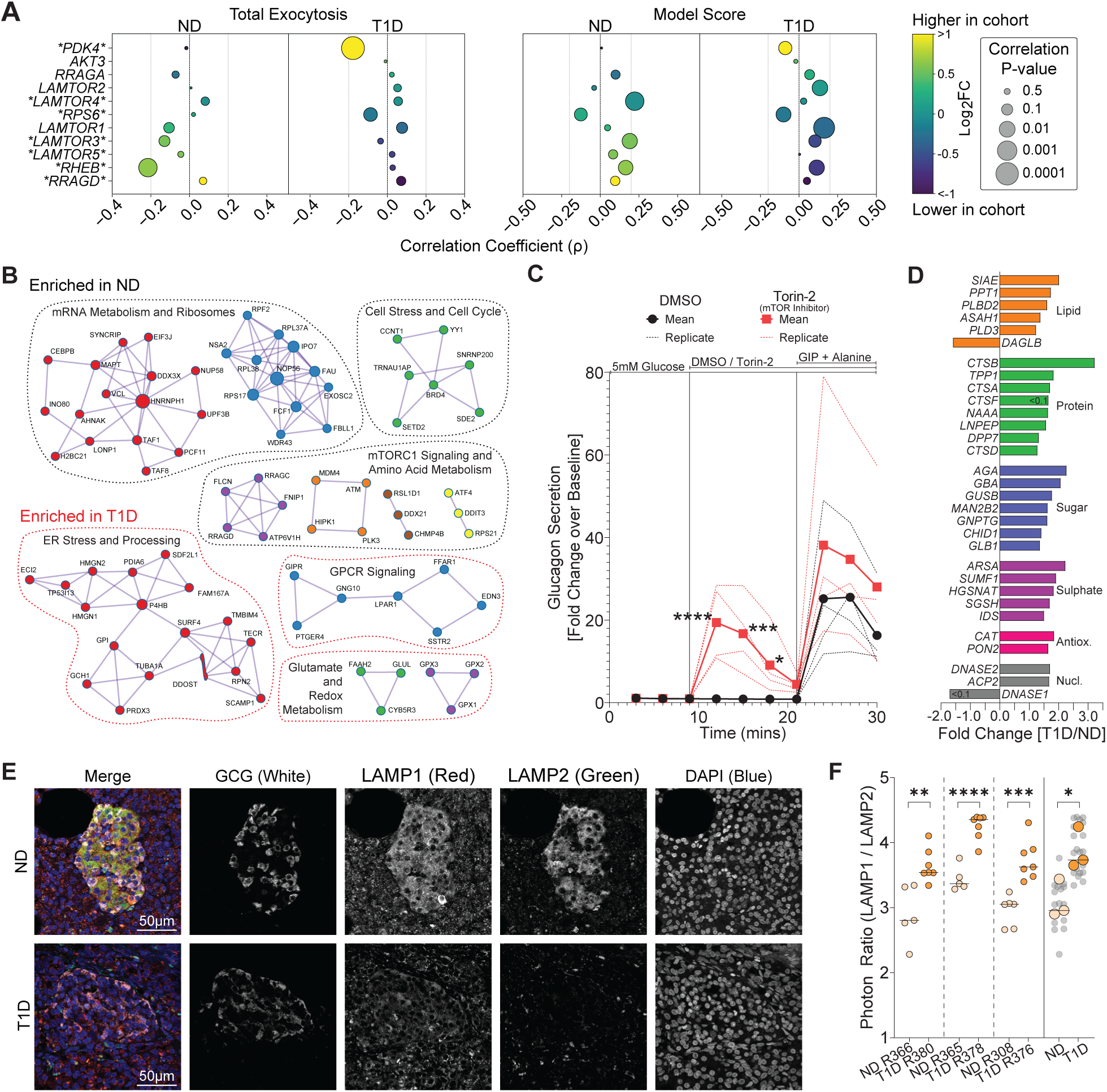
Lysosomal balance in T1D and matched ND donor biopsies, and effects of mTOR inhibition on secretion. (A) Bubble plot of notable transcripts involved with mammalian/mechanistic target of rapamycin complex 1 assembly and activation, their differential expression in T1D vs ND, and their correlative relationship with total exocytosis and model score in α-cells at 5mM glucose. Transcript names flanked by asterisks indicate significant (p<0.05) differential expression. (B) Protein-protein interaction networks of differently expressed genes enriched in ND (black outlines) or T1D (red outlines) α-cells. (C) Glucagon and insulin dynamic secretion profiles in response to mTOR inhibition by Torin-2 (1µM), represented as fold change over baseline from 4 ND donors. (D) Expression of significant (p<0.05, unless indicated) differentially expressed lysosomal associated enzymes in T1D α-cells compared to ND. (E) Representative images of LAMP1 and LAMP2 staining *in situ* from pancreas biopsies of patch-seq donors, attained by immunofluorescence confocal microscopy, and presented in greyscale for better contrast. (F) Quantification of LAMP1:LAMP2 signal ratio in glucagon positive areas. Each data point represents a single field of view of an islet from ND (n = 16, 3 donors) and T1D (n = 21, 3 donors) biopsies. Horizontal line markers indicate median. ^∗^p < 0.05, ^∗∗^p < 0.01, ^∗∗∗^p < 0.001, and ^∗∗∗∗^p < 0.0001 as indicated, using the unpaired t-test (E), or two-way ANOVA (F).

Perturbations to mTORC1 signalling can lead to altered homeostasis of lysosomes, which are implicated in the pathophysiology of diabetes^85,86^. Our DEA analysis shows that lysosomal-associated digestion enzymes^87^ were mostly upregulated in T1D α-cells compared to ND (**Fig 7D**). In α-cells exposed to hyperglycemia, glucagon degradation at LAMP2^+^ lysosomes is reduced, and re-direction to LAMP1^+^ lysosomes is increased, possibly contributing to glucagon hypersecretion^51^. Imaging LAMP1 and LAMP2 (**Fig 7F**) revealed that while both signals appeared lower in T1D, suggestive of altered lysosomal homeostasis, quantification shows the LAMP1/LAMP2 ratio to be higher in T1D α-cells (**Fig 7F**).

## Discussion

Few studies have isolated islets from organ donors with T1D. Single donor studies have suggested a loss of glucose-dependent insulin secretion (despite the presence of some insulin-containing islets) when isolated at^27^ or shortly (∼8 months) after^12^ diagnosis, but perhaps some glucose-responsive insulin release in long-standing T1D^9^. In those single-donor studies, the ability of glucose to suppress glucagon secretion appears lost, although the response to arginine may be preserved^9,12^. More recently, a study on islets of 8 donors with T1D^8^ suggested normal glucose-regulated insulin secretion of remaining β-cells and disrupted glucagon secretion. And islets from 6 donors with T1D (3 of which are included in the present study), had glucagon secretion that was unresponsive to glucose but increased in response to amino acids^88^. In that study, somatostatin secretion was increased from the T1D islets, the antagonism of which also increased glucagon secretion.

We find that the phenotypes of both α- and β-cells from donors with T1D are significantly different than those from ND controls matched for age, sex and BMI. Electrophysiological distinctions can be demonstrated in the activities of key ion channels, exocytotic processes, and aggregate biophysical properties (i.e. the α/β-scores). This suggests a loss of electrophysiologic phenotype in α-cells and surviving β-cells in T1D that likely contributes to dysregulated function, such as the increased α-cell excitability and responsiveness to stimuli we observed. Our results not only capture known transcriptional changes such as the upregulation of antigen presentation and metabolic pathways but also reveal genes and pathways that are likely to play key roles in α-cell dysfunction in T1D by connecting transcriptional changes and biophysical phenotypes. Notably, transcripts linked to exocytotic dysfunction and loss of electrical phenotypes in T1D α-cells are enriched for genes involved in genetic risk of T1D, suggesting a causal relationship between α-cell function and T1D. We also validated some of these identified pathways by linking α-cell dysfunction to increased immune signaling and MHC class I expression, altered expression and localization of key lineage transcription factors, and a role for suppressed mTOR activity and lysosomal balance.

Early impairments in insulin secretion may precede the clinical diagnosis of T1D^4–7^ and have also been reported later in the disease^10–15^. Although it has been suggested some glucose responsiveness in β-cells from a donor with long-standing T1D^9^ and normal insulin responses from surviving T1D β-cells in a larger study^8^, recent studies of live pancreas slices from donors with T1D appear to support the view that the function of remaining β-cells is impaired in T1D^10,14,15^. This is consistent with our demonstration of a loss of electrophysiological phenotype in β-cells from donors with T1D. From a transcriptomic perspective, T1D β-cells appear to have enrichments for glycolysis, and a downregulation for mitochondrial respiration compared to ND, which may suggest a general “stem-like” metabolic phenotype^89^. Indeed, the metabolic interplay between glucose metabolism and mitochondrial respiration observed in stem cell derived β-cells is a driving factor in their questionable secretory behaviour^90,91^. Overall, we provide evidence that the remaining β-cells in T1D have altered electrical and transcriptomic phenotypes compared with matched controls. Due to the limited number of T1D β-cells available however, we focused primarily on a deeper analysis of α-cell phenotypes.

Glucagon secretion is dysregulated in T1D, with a lack of appropriate stimulation by hypoglycemia^20^, and an inappropriately elevated response to a mixed meal^21^. In islets isolated from donors with T1D, low glucose stimulation of glucagon appears impaired, but may be preserved in response to arginine^8,9,12,92^, although this finding has not been universal^15^. Interestingly, glucagon secretion from islets of autoantibody-positive donors appears hyperresponsive to cAMP-raising agents^23^. Consistent with this, we find that islets of T1D donors are hyperresponsive to GIP and alanine. GPCR signaling (including *GIPR* expression) appears upregulated at the transcriptional level in α-cells of T1D donors, where it correlates with inappropriately elevated exocytosis in T1D α-cells. Further, serum levels of GIP have been reported as significantly elevated in T1D patients^93^

The resilience and survival of α-cells in T1D is somewhat enigmatic^94^. One paradigm is that α-cells better handle ER stress and unfolded protein responses^94,95^ in concert with greater resistance to apoptosis via higher expressions of anti-apoptotic markers^96^. Indeed, we see such markers upregulated in T1D, including *BCAP31, BCL2L1, and APIP* (**Suppl Fig 8**). Further, we also find T1D α-cells show an overall upregulation of proteasomal components that correlate with α-score to a greater extent in T1D than in ND, suggesting that in addition to survival, the handling of ER stress and unfolded protein responses may contribute to overall functional behavior, which becomes even more apparent in T1D (**Suppl Fig 8**). Adding to the enigma and further convoluting how they evade autoimmunity in T1D, previous studies show that α-cells not only possess elevated MHC class I^24^ but also seem to preferentially associate with macrophages and T-cells, inadvertently contributing to the killing of neighboring β-cells^97^. Indeed, we see all classical MHC class I markers (*HLA-A/B/C)* and co-molecule *B2M* significantly elevated in T1D α-cells, and correlating with worsening α-scores. Intriguingly, α-cells with the higher expression of immune signaling inhibitors, such as *PIAS1, TOLLIP, and NFKBIA,* trended with improved α-scores. We also observed the compartmentalization of GCG and MHC class I signals in ND α-cells, which appear abrogated in T1D, showing colocalization. This suggests the possibility that the upregulation of antigen presentation may potentially interfere with normal glucagon granule dynamics, contributing to α-cell dysfunction in T1D.

Markers of α-cell identity and immaturity^98^, such as transcription factors *PAX6, NEUROD1* and *ISL1*, have been recently shown as upregulated in T1D α-cells^25^, and our analyses corroborate these findings. We also see that in ND α-cells, higher expression of these markers correlate with a loss of α-score, which may be unsurprising as a subset of α-cells likely possess altered functional behavior due to immaturity, or a transient temporary loss of maturity, which would then drive the expression of these markers^99^. However, the correlation is mostly lost in the T1D α-cells, suggesting a potential breakdown in this feedback mechanism. We questioned why the upregulation for markers that normally drive (re)- maturation still fail to restore α-cell behavior in T1D. Our imaging data reveals that NEUROD1 and ISL1 show a reduction in nuclear localization in T1D. This supports the notion that despite their upregulation, these transcription factors may be insufficient to restore α-cell maturity due to alterations in nuclear accessibility. In fact, immune signaling (and viral infections) can monopolize nuclear transport machinery, leading to a disruption of homeostatic nuclear permeability^81^, and through this, may be inadvertently contributing to a loss of α-cell identity in T1D.

Previous studies of mTOR in α-cells demonstrated that prolonged hyperglycemia leads to sustained mTORC1 activity with elevated glucagon secretion^100^, and that inhibition of mTOR either through α-cell specific RAPTOR knock out^100,101^ or treating with the inhibitor rapamycin^101^ abrogates glucagon secretion. Of note, the RAPTOR knock outs were performed in mouse models and the rapamycin treatments were chronically administered. Further, prolonged rapamycin exposure has been shown to deplete Ca^2+^ stores in the ER and mitochondria, which impairs mitochondrial respiration and reduces secretion^102^. Here we report that under euglycemia (5mM glucose), acute mTOR inhibition by Torin-2, a more potent and selective inhibitor, leads to an increase in glucagon secretion from ND islets. Indeed, mTORC1 components show a positive relationship with α-score, suggesting they are important for function, but show a strong negative correlation with exocytosis in ND that is lost in T1D. When contextualized with the significantly lower expression of mTORC1 components observed in T1D α-cells, such as *RHEB*, multiple *LAMTORs,* and *RRAGD*, it suggests that at least at 5mM glucose, these alterations to mTOR signaling may be contributing to the observed secretory dysfunction. This may in fact be linked to MHC class I presentation disrupting glucagon granule trafficking mentioned earlier, as the lack of mTOR signaling can induce MHC class I overexpression^103^. The alterations to mTORC1 in the T1D α-cell may also hold implications with lysosomal balance. Previous studies in αTC1-6 cells, a cell line model for α-cells, demonstrate that under chronic hyperglycemia, glucagon granules are mis-routed from degradative LAMP2^+^ lysosomal compartments and directed towards LAMP1^+^ secretory granules, contributing to glucagon hypersecretion^51,77^. Although a full characterization of the mechanism is outside the scope of this publication, it appears clear that a contributor to T1D α-cell dysfunction may lie in a potential concerted interplay amongst alterations to the proteasome, mTORC1 signaling, and lysosomal balance. Indeed, a common currency linking these cellular components together are free amino acid pools, and altered profiles of free plasma amino acids is a well-known characteristic of T1D^104,105^.

Finally, while genetic risk for T1D is strongly linked to T-cell activity, T1D risk variants have also been implicated in β-cells and other pancreatic cell types such as acinar and ductal cells^106^. Here we find novel evidence that genes up-regulated in T1D α-cells are linked to T1D risk variants, supporting a potential causal link between α-cells and T1D. Perhaps more interestingly, we find enrichment of T1D-associated genes within transcriptional anti-correlates to α-score measures in T1D α-cells, and a depletion within positive correlates of α-score, suggesting that α-cells with a more severe disruption of electrophysiologic phenotype (i.e. a loss of a-score) have higher expression of transcripts associated with T1D risk, while cells with preserved α-scores express lower levels of these. Thus, electrophysiological dysfunction in α-cells may increase with the risk of T1D. Indeed, a key contributor to the enrichment was MHC class 1 component genes which are major factors in T1D genetic risk. Altogether, this suggests links between α-cell dysfunction and genetic susceptibility to T1D.

In summary, we reveal significant functional and molecular differences in both α- and β-cells from donors with T1D consistent with altered regulation of glucagon and insulin secretion. Transcriptomic analyses link these functional alterations to dysregulated immune signaling, antigen presentation (notably MHC class I overexpression), and disruptions in key cellular pathways, including mTOR signaling and lysosomal balance. Markers of α-cell lineage such as *NEUROD1* and *ISL1* are upregulated in T1D, yet their expected regulatory feedback mechanisms appear impaired, likely due to altered nuclear localization of these transcription factors. Some functional alterations in T1D α-cells may be directly linked to genetic disease risk, reinforcing the idea that genetic susceptibility extends beyond immune and β-cell compartments. Overall, our findings provide a mechanistic link between transcriptional, electrophysiological, and functional disruptions in pancreatic islet cells in T1D, highlighting their potential contribution to disease pathophysiology.

### Limitations

We have performed due diligence to only include control donors that match the T1D cohort when comparing our findings. However, we do note that the sex of both cohorts is skewed towards males, which is solely influenced by donor availability. While the present study links functional and molecular phenotypes of α-cells in T1D, we recognize that dissociated islet cells may behave differently than cells within intact tissues. There may be several reasons for this including loss of paracrine signaling, cell-cell interactions, and other structural impacts of the microenvironment. This may be addressed in part in ongoing studies of live pancreas slices^10,14,15,92^, although a direct connection between molecular profiles and *in situ* (dys)function remains a challenge. We have performed validation studies through tissue sections, and glucagon secretion measurements from intact islets, which provides some level of confidence that key single-cell findings hold within the intact tissue. Additionally, our ability to detect well-established pathways in T1D, such as immune signalling pathways, also provides some reassurance regarding the validity of results.

Protein level validation and quantification would ideally be performed using a more direct biochemical approach such as immunoprecipitations and western blotting; however, due to the limited availability of T1D donor tissue, and low abundance of islets in pancreata, protein yield would be insufficient for such methods. Further, animal models of T1D fail to sufficiently capture the human T1D phenotype^107^. Thus, we imaged biopsies to gain *in situ* protein information, using ratiometric quantification or colocalization approaches which self-normalize to the sample being imaged. We avoided direct comparisons of photon signals between samples, which can be influenced by FFPE sample dimensions, preservation age, tissue density, sectioning variation, and other technical factors.

While we show that MHC class I localization with glucagon and nuclear accessibility are altered in the T1D α-cells, and this could be linked to the upregulation of immune signalling pathways, a fuller understanding of the direct mechanisms involved in these alterations would require additional characterization of endo-lysosomal compartments and their trafficking machinery and assessing nuclear pore complex composition, respectively. Further, it is uncertain whether enhancing nuclear permeability for α-cell transcription factors will restore normal secretory behaviour in T1D. Finally, although mTOR inhibition in ND islets results in an increase in glucagon secretion, it does not fully mimic the mTORC1 phenotype in T1D α-cells, where there is an abrogation in mTORC1 component expression, which will likely result in insufficient assembly of the complex. Thus, agonism of mTOR may not rescue α-cell secretory behaviour in T1D, unless the complex successfully assembles first.

## Materials and Methods

### Human islets from donors with and without T1D

Majority of human islets used in this study were isolated at the Alberta Diabetes Institute IsletCore, isolated most of the islets for this study, using established protocols^108^. Detailed information on all donors, quality control, and islet cell phenotyping assays are described at www.HumanIslets.com^29^. Additional T1D samples were from the nPOD islet isolation program, the Human Pancreas Analysis Program (HPAP)^109^, and from R. Bottino. Donors included in this study are listed in **Supplementary Table 1**. All studies were performed with approval of the Human Research Ethics Board at the University of Alberta (Pro00013094; Pro00001754). All donors’ families gave informed consent for the use of pancreatic tissue in research.

The study cohort included all patch-seq cells obtained from donors with diagnosed T1D, resulting in a range of age from 13 to 44 years, Body Mass Index (BMI) of 17.0 to 29.4 kg/m^2^, and Cold Ischemia Time (CIT) of 2 to 24.5 hours. For the control cohort, we used patch-seq cells identified as α- or β-cells from donors not diagnosed with diabetes (ND), whose age, BMI, and CIT ranged between 12 to 46 years, 16.4 to 27.5 kg/m^2^, and 2 to 23 hours respectively. We excluded ND patch-seq data from donors with characteristics outside these ranges to maintain similarity with our T1D donor cohort, and help minimize potential effects from age, BMI, and CIT on subsequent analyses.

### Hormone secretion assays

To measure dynamic glucagon and insulin secretion, 35 islets per group were pre-incubated for 30-45 min in KRB buffer containing (in mM): 140 NaCl, 3.6 KCl, 2.6 CaCl_2_, 0.5 NaH_2_PO_4_, 0.5 MgSO_4_, 5 HEPES, 2 NaHCO_3_ and 0.5 mg/ml essentially fatty acid free BSA and glucose as indicated using the Biorep Peri4 perifuison system (Biorep). Islets are perifused with KRB supplemented with glucose concentrations, GIP (Anaspec) and the amino acid alanine (Sigma) and mTOR drugs as indicated in figures. Total content was obtained by lysing the cells with acid ethanol. Samples are treated with 5ug/ml Aprotinin (Sigma) to prevent degradation. All the samples were collected and stored at -20 °C for ELISA of glucagon (MSD; K1515YK-2) and insulin (Alpco; 80-INSHU-CH01).

### Electrophysiology

*<u>K_ATP_ channel activity</u>* was measured in whole-cell configuration upon washout of intracellular ATP. Whole-cell currents were recorded in response to voltage steps between −60 and −80 mV from a holding potential of −70 mV. The bath solution contained 138 mM NaCl, 5.6 mM KCl, 1.2 mM MgCl_2_, 2.6 mM CaCl_2_, 5 mM HEPES, and 5 mM glucose (pH 7.4). The pipette solution contained 125 mM KCl, 30 mM KOH, 1 mM MgCl_2_, 10 mM EGTA, 5 mM HEPES, 0.3 mM Mg-ATP, and 0.3 mM K-ADP (pH 7.15).

*<u>Action potentials</u>* were measured in current-clamp mode of the perforated patch-clamp configuration. The bath solution contained (in mM): 140 NaCl, 3.6 KCl, 1.5 CaCl_2_, 0.5 MgSO_4_, 10 HEPES, 0.5 NaH_2_PO_4_, 5 NaHCO_3_, and indicated concentrations of glucose (pH 7.3 with NaOH). The patch pipettes were filled with (in mM): 76 K_2_SO_4_, 10 KCl, 10 NaCl, 1 MgCl_2_ and 5 HEPES (pH 7.25 with KOH), and backfilled with 0.24 mg/ml amphotericin B (Sigma, cat# a9528). Quality control was assessed by the stability of seal (>10 GOhm) and access resistance.

<u>Cell identity</u> was determined by immunostaining for glucagon (primary antibody: monoclonal anti-glucagon antibody produced in mouse, 1:200, Sigma cat# G2654; secondary antibody: 1:200, Alexa Fluor 594 goat anti-mouse, Thermofisher cat# A-11032) and insulin (primary antibody: 1:200, DAKO/Agilent IR00261-2; secondary antibody: 1:200, Alex Fluor 488, Thermofisher cat# A-11073). Data were analyzed using FitMaster (HEKA Electronics) and Prism 10 (GraphPad Software Inc., San Diego, CA).

*<u>Electrophysiology for patch-seq</u>* was performed as previously^17,26^. In brief, hand-picked islets were dissociated to single cells using StemPro accutase (Thermo Fisher Scientific, Cat#A11105-01) and plated on 35-mm cell culture dishes for patch-clamp and patch-seq as previously^17,26^. Single cells were cultured in low glucose (5.5 mmol/L) DMEM with L-glutamine, 110 mg/L sodium pyruvate, 10% FBS, and 100 U/mL penicillin/ streptomycin for 1-3 days before patch-clamp in the whole-cell or perforated patch-clamp configuration in a heated bath at 32-35°C using pipettes with resistances of 4-5 MOhm after fire polishing. Whole-cell capacitance responses as a measure of exocytosis were recorded with the Sine+DC lock-in function of a HEKA EPC10 amplifier and PatchMaster software (HEKA Electronics, Germany) in response to ten 500 ms depolarizations to 0 mV from a holding potential of -70 mV, and Ca^2+^ and Na^+^ currents were activated by the membrane potentials ranging from -70 to -10 mV. Responses were measured 1-2 minutes after obtaining the whole-cell configuration. The bath solution contained (in mM): 118 NaCl, 20 tetraethylammonium-Cl, 5.6 KCl, 1.2 MgCl_2_, 2.6 CaCl_2_, 5 HEPES, and 5 glucose (pH 7.4 with NaOH). Patch pipettes were filled with (in mM): 125 Cs-glutamate, 10 CsCl, 10 NaCl, 1 MgCl_2_, 0.05 EGTA, 5 HEPES, 0.1 cAMP and 3 MgATP (pH 7.15 with CsOH).

### Single-cell RNA sequencing and analysis

*<u>Single-cell RNA sequencing</u>* was performed on patched cells collected using a separate wide-bore collection pipette (0.2-0.5 MOhm) containing lysis buffer (10% Triton, Sigma-Aldrich, #93443; ribonuclease inhibitor 1:40, Clontech, #2313A), ERCC RNA spike-in mix (1:600000; ThermoFisher, #4456740), 10 mM dNTP, and 100 μM dT (3’-AAGCAGTGGTATCAACGCAGAGTACTTTTTTTTTTTTT TTTTTTTTTTTTTTTTTVN-5′) and then stored at -80°C in PCR tubes. cDNA and sequencing libraries were generated via an adapted SmartSeq-2 protocol for patch-seq plates^17,26,110^. Library generation was via amplification of the cDNA by Tn5 tagmentation, and sequenced on the NovaSeq (Illumina) using paired-end reads (100bp) to an average depth of 1 million reads per cell. Alignment was against the human genome (GRCh38 with supplemental ERCC sequences) using STAR^111^, with gene counts determined via htseq-count (intersection-nonempty) using GTF annotation with Ensembl 89 release genes^112^.

*<u>Cell typing of patch-seq cells</u>* was performed in ScanPy using the scRNA-seq data of all ND and T1D patch-seq cells available, with gene expression normalized to 10,000 (i.e. 1e4) and transformed using natural logarithm. Variations in total counts and percent mitochondrial counts were regressed out prior to Uniform Manifold Approximation and Projection (UMAP) generation and Leiden clustering. Clusters with the expression of Insulin (INS) and Islet Amyloid Polypeptide (IAPP) were identified as β-cells, Glucagon (GCG) as α-cells, Somatostatin (SST) as Delta cells, Pancreatic Polypeptide (PPY) as Gamma cells, Serine Protease 1 and 2 (PRSS1, PRSS2) as Acinar cells, and Ghrelin (GHRL) as Epsilon cells (of which there were no distinct clusters present).

*<u>Differential expression analysis (DEA)</u>* was performed using the Wilcoxon rank-sum method to compare the transcriptomes of α- and (surviving) β-cells between the T1D and matched control ND donors, similar to the approach outlined above, except with total counts set to 1,000,000 (1e6) to maintain consistency with prior works when performing subsequent analyses^17,26^. Following UMAP generation, we overlaid additional metadata, including donor ID, cell type, and donor characteristics such as HbA1c, diabetes status, BMI, Years with Diabetes, Age, and Sex. We also overlaid select corresponding electrical data, including cell size, total exocytosis, and sodium currents. To identify enriched pathways, we used the WEB-based GEne SeT AnaLysis Toolkit^113,114^ (WebGestalt) to perform Gene Set Enrichment Analyses (GSEA) on the transcript’s DEA score, with a False Discovery Rate (FDR) threshold of 0.1, and Weighted Set Cover for redundancy reduction.

### Patch-seq Analysis

For *<u>electrophysiological fingerprinting</u>*we integrated and scored electrical data using a gradient boosted decision tree-based ensemble machine learning algorithm as described previously^26^. In brief, we used Extreme Gradient Boosting (XGBoost v1.6.2) in a Python v3.7.11 framework to classify cells as α- or β-cells based solely on electrophysiology data, without *a priori* knowledge of cell identity, and provide a numerical metric of confidence. The model used electrical data from the ND patch-seq cohort, and from cells of ND donors of similar age, BMI, and CIT, identified by immunostaining (α-cells: 248 and 9, β-cells: 104 and 186 respectively). Electrophysiological data included were cell size (pF), normalized total capacitance (fF/pF), normalized first depolarization capacitance (fF/pF), normalized late depolarized capacitance (fF/pF), normalized early peak calcium current amplitude (pA/pF), normalized late calcium current amplitude (pA/pF), calcium integral normalized to cell size (pC/pF) and normalized peak sodium current amplitude (pA/pF). Fine tuning was performed to attain a predetermined minimum accuracy of 75%, with early stopping of 100 iterations and AUCPR as the evaluation metric. Confusion matrices were generated using scikit-learn (1.0.2) to assess training performance, and the feature importance graph via XGBoost to ensure that electrical parameters provided were used in decision making without disproportionate reliance on a single parameter(s)^17^. When applied to the ND and T1D patch-seq cell cohorts the predicted probability scores were taken as the numerical metric to assess fit to α-cell (α- probability = 1.0) and β-cell (α-probability = 0.0) canonical electrical behavior. To determine whether donor characteristics significantly trended with model scores and electrophysiology, Ordinary Least Squared (OLS) regression was employed using statsmodel (v0.12.2) and sklearn (v1.0.2).

*<u>Spearman’s rank correlations</u>* were performed using SciPy between transcript expression and electrophysiology for transcripts detected in at least 50% of the cohort. Cells with undetectable expressions for a transcript, were excluded from only that specific correlation. We employed bootstrapping (1000 iterations) and performed GSEA using the mean bootstrapped transcript correlation coefficients as weighting, or Over Representation Analysis in WebGestalt.

*<u>α-cell protein-protein interaction (PPI) networks</u>* were generated using Metascape^115^. Briefly, a list of genes significantly expressed in ND or T1D α-cells (log2FC > or < 1, p-value < 0.05 were uploaded to Metascape web browser tool for pathways enrichment analysis and the Metascape MCODE network function was used to create PPI networks. Only nodes with 3 or more connections are shown.

*<u>Enrichment of genes associated with T1D risk alleles</u>* was performed by compiling gene sets associated with T1D risk from multiple sources. First, we identified genes from the GeneSifter tool on the T1D Knowledge Portal^116^ (T1DKP; https://t1d.hugeamp.org/) with the following filters: TPM ≥ 1 in the pancreas and T1D Human Genetic Evidence^117^ (HuGE) score ≥ 3. Second, we identified genes with pancreatic islet expression QTLs from the InsPIRE study^118^ for T1D-associated variants (p<5x10^-8^) from a published GWAS^106^. We defined strict and permissive sets of islet eQTLs which were based on genome wide significance (2x10^-6^) or more nominal significance (5x10^-3^), respectively. We then used Fisher’s Exact Test to calculate the enrichment of each set of T1D-associated genes in genes enriched in ND and T1D α-cells. Briefly, we compared the number of T1D risk genes associated with ND or T1D α-cells with all genes tested in these association tests and calculated two-sided p-values for enrichment.

### Pancreatic Biopsy Imaging

*<u>Slide samples</u>* were prepared from formalin fixed paraffin embedded (FFPE) pancreas biopsies of ND and T1D pancreata, sectioned at 4μm by the Alberta Diabetes Institute HistoCore. Samples were heated at 58°C for 20mins, followed by deparaffinization and rehydration using graded xylene → ethanol → water washes. Heat induced epitope unmasking was performed using antigen retrieval buffer (10mM citrate, 0.05% Tween-20, pH 6.0) and a microwave oven at 680W for 16 mins. For the following steps, three washes with PBS were performed prior to moving to the next step: Permeabilization was performed with 0.1% Triton X-100 in PBS for 10mins and blocking with 20% goat serum in PBS for 30 mins at room temperature. Primary antibodies, except for anti-glucagon, were diluted in 4% goat serum in PBS and used at concentrations between 1-2μg/mL, applied as a 175μL droplet directly to the FFPE section, and incubated in the absence of light, at 4°C overnight, in a humidified chamber. Secondary antibodies were diluted to 5μg/mL in 4% goat serum and were applied as a 175 μL droplet directly to the FFPE section and incubated in the absence of light for 1 hour at room temperature, in the humidified chamber. Anti-glucagon primary was applied at 1μg/mL for 30mins at room temperature, followed by its secondary and DAPI for 20mins. Samples were mounted using 50μL of Prolong Glass with a 170mm high precision (1.5H) coverslip and cured overnight at room temperature in the dark.

*<u>Image acquisition</u>* was performed on the Leica STELLARIS 8 DMI8 platform, with a 63x 1.4 numerical aperture oil immersion lens (refractive index 1.518). DAPI was excited with a 405nM diode laser, and secondaries were excited with the white light laser using the LAS X (5.2.2) acquisition software’s dye assist specifications, which gated emission spectra to eliminate channel cross talk. All detectors were configured for 8-bit photon counting. Excitation parameters were kept identical when imaging between samples. LAS X navigator was used to locate islets via the glucagon signal, after which, multi-channel 3D images were acquired at 1024x1024 resolution, using Z-stacks, with 0.2µM steps. Each channel was excited and captured independently prior to moving to the next Z-stack. Deconvolution was performed using LAS X’s Lightning Deconvolution.

*<u>Image analysis</u>* was performed in Fiji (v1.54k.). The Lightning Deconvolution decision heatmap for the glucagon signal indicated the boundaries of α-cells, allowing for quantification of signals specific to glucagon areas in the islet. Ratiometric signal quantification was performed on raw signals from “summation” hyperstacked images. For measuring % nuclear signal, regions of interests were user defined using the DAPI signal to define the cell’s nuclei, and the glucagon signal to identify its associated cytoplasm. 3D co-localization analyses used Fiji’s Coloc2 plugin across Z-stacks with the Pearson correlation coefficient method and were performed with Lighting deconvoluted images.

### Figure Generation

Figures involving the analysis of transcriptomic data were generated either using a combination of Scanpy (1.9.3), Seaborn (0.12.2) and Matplotlib (3.4.2). Graphs comparing electrophysiology data were generated in GraphPad Prism 10 using the Box and Whisker preset, with whiskers indicating the 10-90 percentile of data. Microscopy images were generated in Fiji (v1.54k) from the raw LAS X imaging files acquired on the STELLARIS platform. Layouts and graphics were designed in Adobe Illustrator (28.5).

### Data Availability

Raw sequencing reads are available in the NCBI Gene Expression Omnibus (GEO), and Sequence Read Archive (SRA) under accession numbers GSE124742, GSE164875, and GSE270484. Patch-seq and other human islet data are available for analysis on the HumanIslets.com resource.

## Acknowledgements

The University of Alberta acknowledges that we are located on Treaty 6 territory, and respects the histories, languages, and cultures of First Nations, Métis, Inuit, and all First Peoples of Canada, whose presence continues to enrich our vibrant community.

We thank all families and donors for the generous gifts in support of diabetes and transplantation research and Give Life Alberta (Alberta Health Services), the Trillium Gift of Life Network (TGLN), BC Transplant, Quebec Transplant, and other Canadian organ procurement organizations for their efforts procuring pancreas for research. We also thank Dr. Rita Bottino (Director Islet Programs, Imagine Islet Center – Imagine Pharma, Pittsburgh PA), the nPOD Islet Isolation Pilot Program led by Dr. Clayton Mathews (University of Florida, Gainesville, FL), and the Human Pancreas Analysis Program (HPAP; https://hpap.pmacs.upenn.edu) for some additional T1D samples.

We thank the Alberta Diabetes Institute HistoCore (Lynette Elder, Dr. Gregory Korbutt) for assistance with biopsy sectioning, and the University of Alberta Cell Imaging Core for their microscopy facilities (Kiera Smith, Dr. Hilmar Strickfaden).

## Funding

This work was supported by grants from the Canadian Institutes of Health Research (MacDonald - 186226), Breakthrough T1D (MacDonald - 2-SRA-2019-698-S-B), and the National Institutes of Health (MacDonald - DK120447; Gaulton - DK105554, DK138512, HG012059). Some data collection was supported by the Human Pancreas Analysis Program (HPAP-RRID:SCR_016202), a Human Islet Research Network (RRID:SCR_014393) consortium (UC4-DK-112217, U01-DK-123594, UC4-DK-112232, and U01-DK-123716). TdS was supported by the Alberta-Helmholtz Diabetes Research School, the Alberta Innovates Scholarship in Data-Enabled Innovation, and the Sir Fredrik Banting and Dr. Charles Best Canada Graduate Scholarship. HMM was supported by the DT O’Connor Scholar in Genetics award from the University of California San Diego. JCS was supported by the Knut and Alice Wallenberg Foundation (Wallenberg Molecular Medicine Fellow), the Swedish Research Council (grant 2021-05109), and the Erling Persson Foundation (Swedish Foundations’ Starting Grant). KJG holds an endowed chair from the University of California San Diego. PEM holds the Canada Research Chair in Islet Biology.

## Author contributions

P.E.M. and S.Q. conceived the study, with contributions from T.d.S. to experimental design and interpretation. X.-Q.D. performed the electrophysiology, with sequencing performed by J.C.S., R.J., N.F.N, A.D., and M.T.. The scRNA-seq analysis was performed by T.d.S. with contributions from J.D.E., C.E.E., J.C.S. and J.X. GWAS analysis were performed by H.M. and K.G. Electrophysiological fingerprinting and model design were performed by T.d.S. and P.E.M., with contributions from X.-Q.D., A.B., and J.C.S. Perifusion experiments were performed by A.F.S and T.d.S. Islet isolation and biopsies were performed by J.G.L., N.S., A.B., and J.E.M.F. Slide preparation, staining, and microscopy was performed by T.d.S. The protein-protein network analysis was performed by R.A.D.. T.d.S. and P.E.M designed graphics. T.d.S and P.E.M wrote the manuscript, and all authors contributed to editing the final version. P.E.M. acts as the guarantor of this work and is responsible for data access.

## Notes

https://www.humanislets.com

### Competing Interest Statement

The authors have declared no competing interest.

