## Supplemental Figures for "Linking molecular pathways and islet cell dysfunction in human type 1 diabetes"


#### Highlights

- Surviving β-cells in T1D show disrupted electrical function linked to metabolic reprogramming and immune stress.
- Transcripts associated with α-cell dysfunction are enriched in genetic risk alleles for T1D.
- Upregulated MHC class I and impaired nuclear localization of key transcription factors associate with α-cell dysfunction in T1D.
- T1D α-cells exhibit increased hyper-activity, lysosomal imbalance and impaired mTORC1 signaling, which promotes dysregulated glucagon secretion.


*Correspondence to:
Patrick MacDonald
Alberta Diabetes Institute
University of Alberta
LKS Centre, Rm. 6-126
Edmonton, AB
Canada, T6G2R3

www.bcell.org

***Supplemental Figure 1: Cell typing of ND and T1D patch-seq data***

(A) UMAP of ND (n=1435, 46 donors) and T1D (n=990, 10 donors) patch-seq cells with Leiden cluster overlays, and a dot plot of their expression for islet cell type markers.

(B) UMAP from A with islet cell type marker expression overlays.

(C) UMAP from A with clusters identified for cell type, and a dot plot of their expression for islet cell type markers.

***Supplemental Figure 2: UMAP of cell typed ND and T1D patch-seq cells with donor meta data overlays***

Overlays of donor information include age (years), body mass index ($kg/m^2$; BMI), glycated hemoglobin A1c (%; HbA1c), sex, diabetes status, duration of diabetes (years), and donor identification codes (ID).

***Supplemental Figure 3: Comparison between the T1D and matched ND donors from which patch-seq data was analyzed***

For the 9 T1D donors included in this study, we matched them against 17 ND control donors based on the T1D donor characteristics, which included an age range between 12 to 46 years, body mass index (BMI) between 16.4 and 27.5 $kg/m^2$, and cold ischemia time of 2 to 23 hours. One T1D donor (R296) was dropped from this study due to suspiciously elevated insulin content, suggesting a possible *erratum* in reported diabetes status.

***Supplemental Figure 4: UMAP of patch-seq data from T1D and matched control ND donors.***

(A) UMAP of T1D (n=692, 9 donors) and control ND (n=375, 17 donors) patch-seq cells, with overlays of cell type, and a dot plot of their expression for islet cell type markers.

(B) UMAP from A with islet cell type marker expression overlays.

(C) UMAP from A with diabetes status overlay, and a dot plot of their expression for islet cell type markers.

***Supplemental Figure 5: UMAP of patch-seq data from T1D and matched control ND donors with donor characteristics and electrophysiology***

Overlays of donor information include donor identification codes (ID), age (years), body mass index ($kg/m^2$; BMI), glycated hemoglobin A1c (%; HbA1c), sex, diabetes status, duration of diabetes (years), and electrophysiology recordings of cell size (pF), total exocytosis (fF/pF), and peak $Na^+$ current (pA/pF).

***Supplemental Figure 6: Impact of donor characteristics on electrophysiology and α-score***

Ordinary least squares (OLS) regression of indicated electrophysiology and model score values against donor characteristics. OLS analysis was repeated for the T1D data with diabetes duration as an additional variable, which is not applicable for the ND data.

***Supplemental Figure 7: Cell typing of ND and T1D patch-seq data***

(A) Comparison of the electrophysiology and model scoring at differing glucose concentrations obtained from patch-seq α-cells during whole-cell patch-clamp. At 1mM glucose, ND (13 donors) and T1D (4 donors) n-values for cell size = 156, 81; total exocytosis = 154, 81; early exocytosis = 154, 80; late exocytosis = 157, 80; peak $Na^+$ current = 154, 78; early $Ca^{2+}$ current = 156, 80; late $Ca^{2+}$ current = 154,80; $Ca^{2+}$ charge entry = 149, 75; exocytosis normalized to $Ca^{2+}$ = 136,74. At

10mM glucose, ND (10 donors) and T1D (4 donors) n-values for cell size = 85, 80; total exocytosis = 84, 80; early exocytosis = 84, 80; late exocytosis = 84, 80; peak $Na^+$ current = 84, 81; early $Ca^{2+}$ current = 84, 80; late $Ca^{2+}$ current = 83, 79; $Ca^{2+}$ charge entry = 77, 70; exocytosis normalized to $Ca^{2+}$ = 68,69.

(B) α-scores of electrophysiology recorded at 5mM, 1mM, and 10mM from patch-seq alpha cells of ND matched controls (17, 13, 10 donors respectively) and T1D (9, 4, 4 donors respectively). ND and T1D n-values for score at 5mM = 248, 596; 1mM = 158 ,82; 10mM = 85, 81.

(C) Cumulative frequency graph of α-scores of patch-seq α-cells patched at 1mM, 5mM, and 10mM glucose.

*p < 0.05, **p < 0.01, ***p < 0.001, and ****p < 0.0001 as indicated using the two-tailed non-parametric Mann-Whitney test in (A) and using the non-parametric Kruskal-Wallis test with Dunn's correction (B). Outliers were determined based on |z-score| >3 and excluded from comparison and statistical analysis (A and B).

***Supplemental Figure 8: Integrated analysis of notable transcript hits from pathways that may contribute to α-cell dysfunction***

Bubble plot combining differential expression analysis, and correlations of transcript expression with either total exocytosis or α-score in α-cells patched at 5mM glucose. Hits are categorized based on their physiological roles. Transcript names flanked by asterisks indicate significant (p<0.05) differential expression.

***Supplemental Figure 9: Linear regression of nuclear ISL1 and NEUROD1 signals***

****p < 0.0001 as indicated

### Supplemental Figure 1

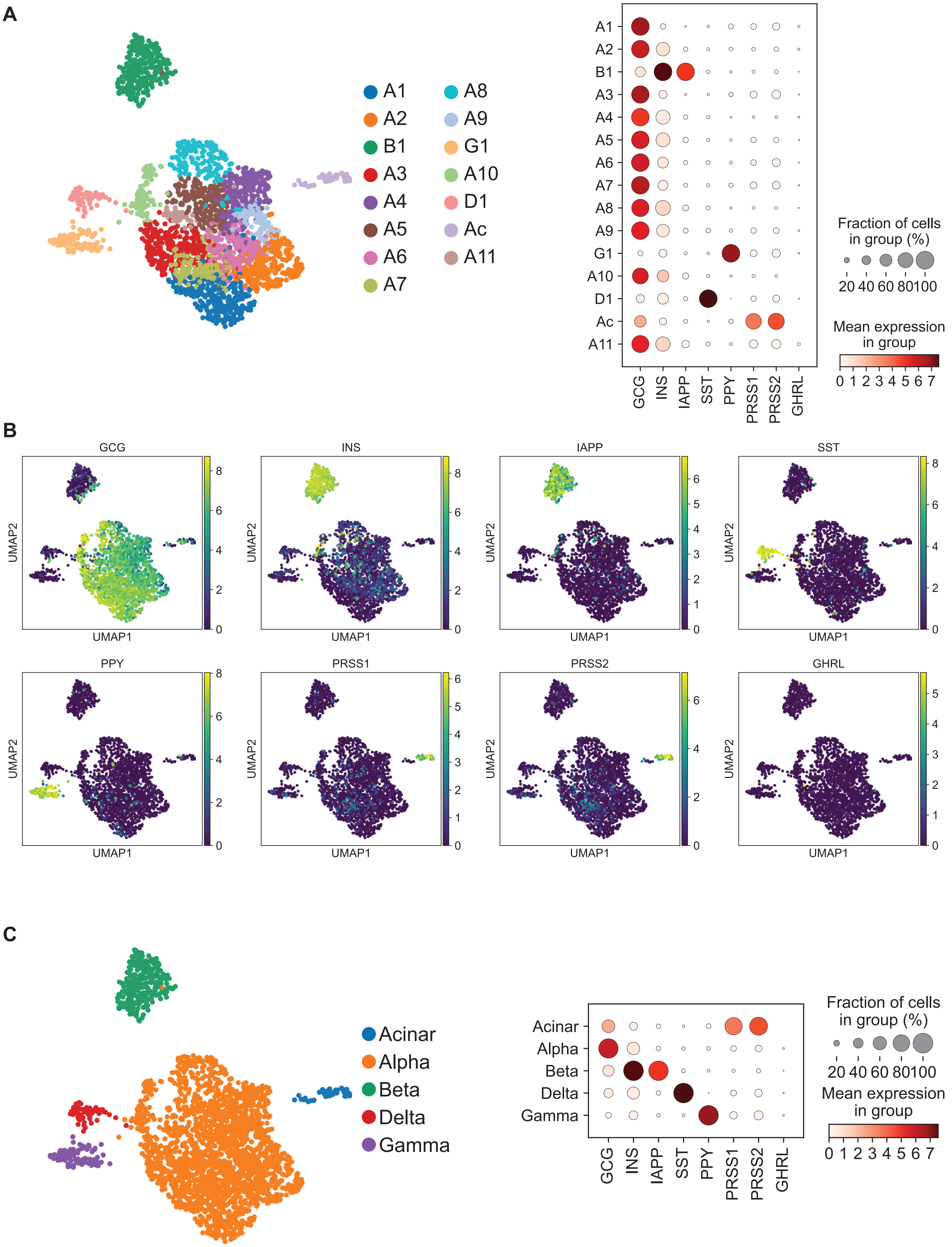

### Supplemental Figure 2

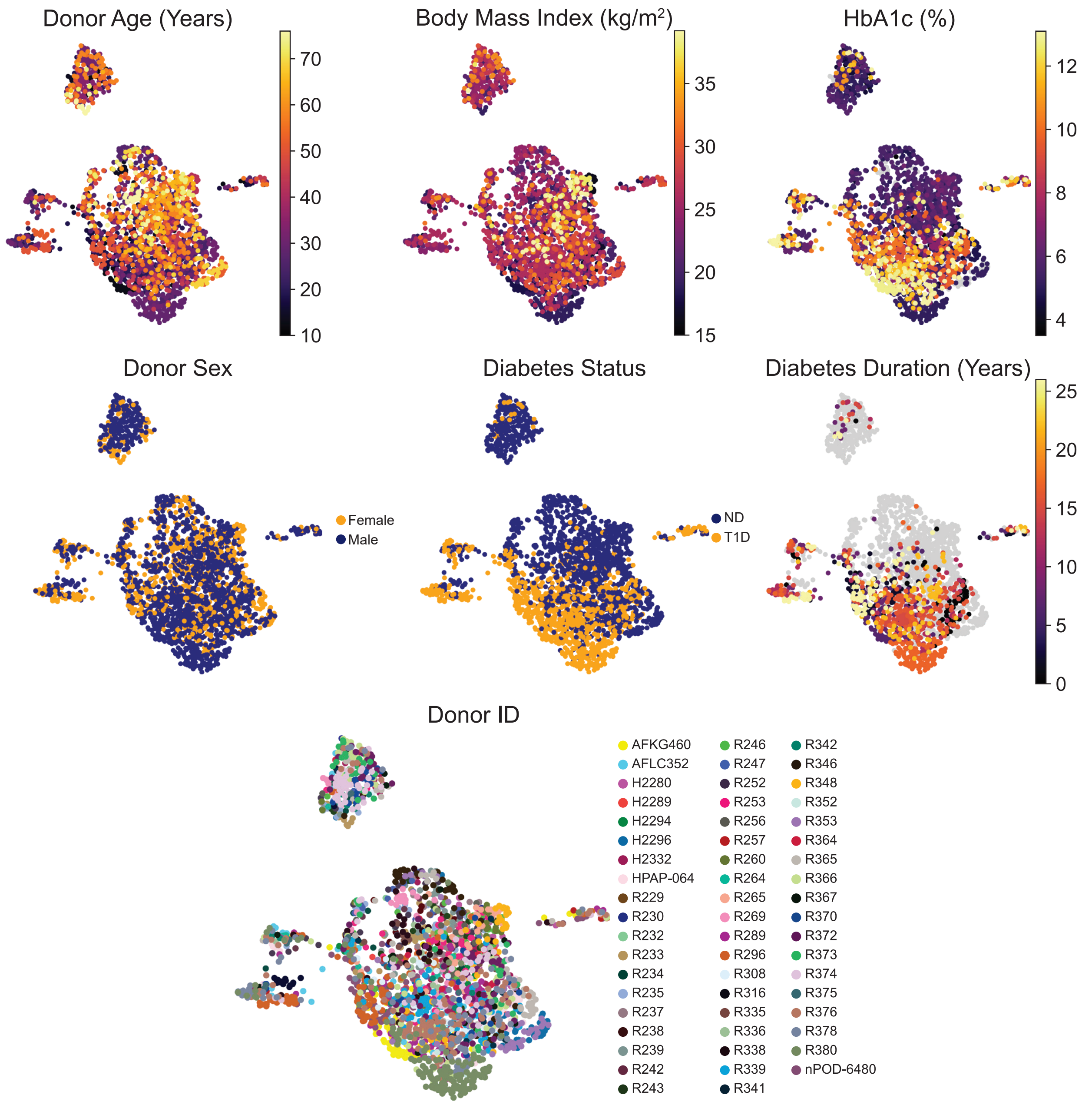

#### Supplemental Figure 3

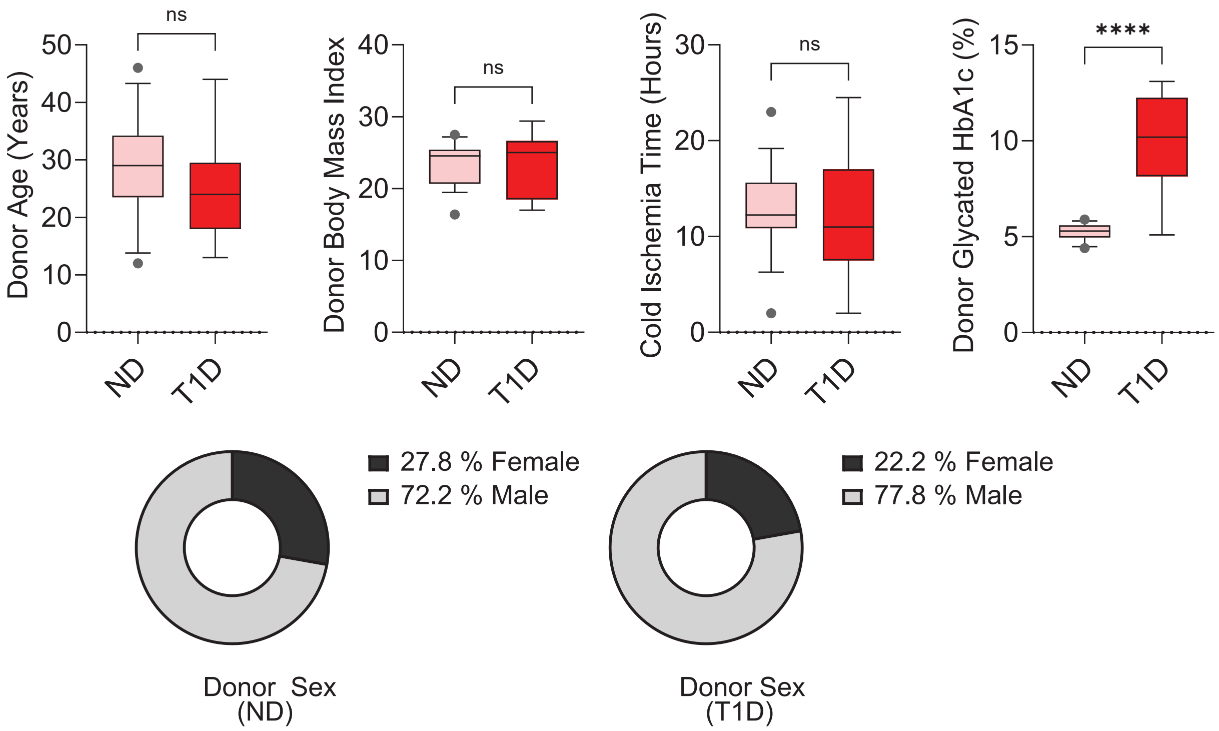

### Supplemental Figure 4

A

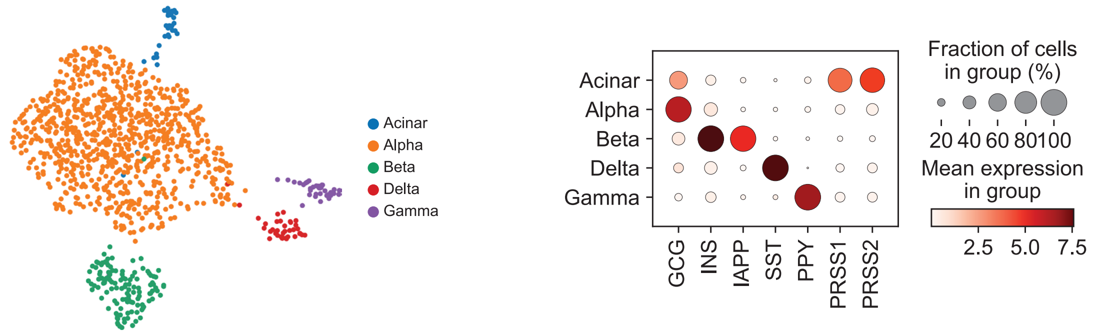


B

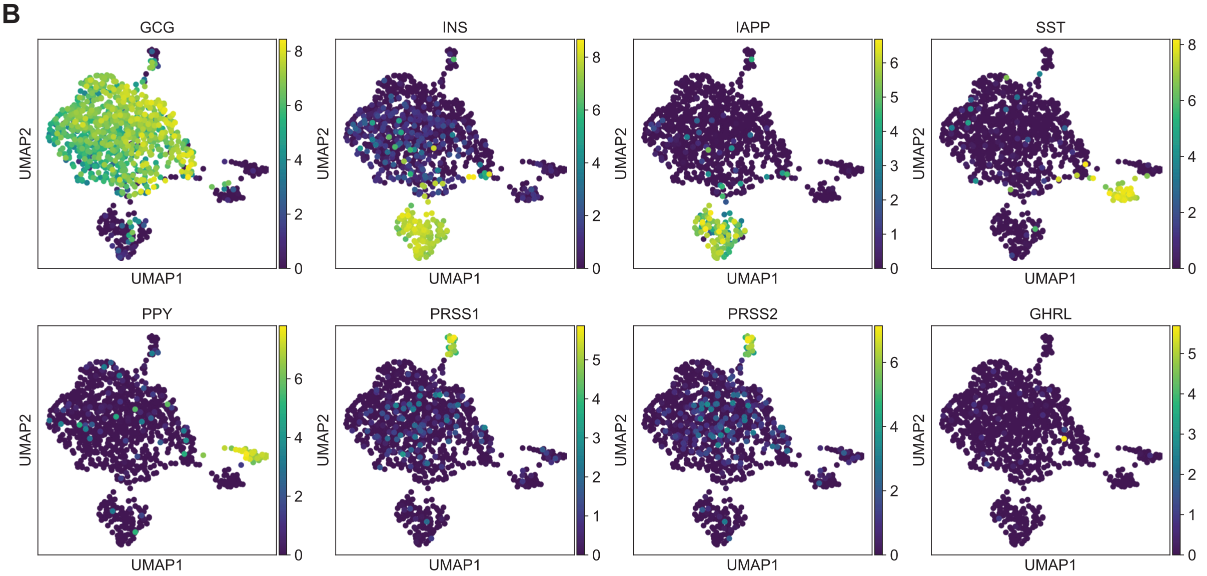


C

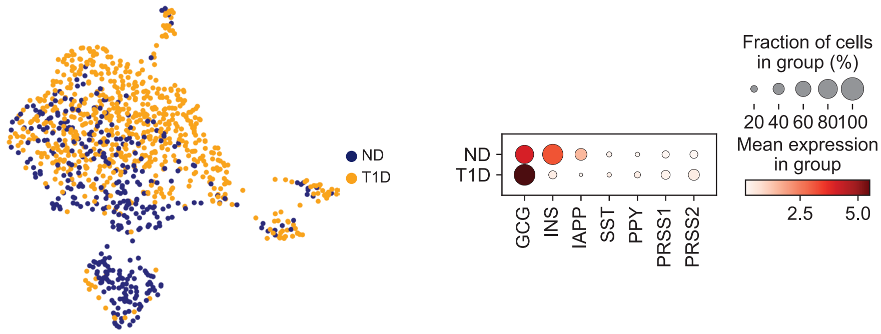

### Supplemental Figure 5

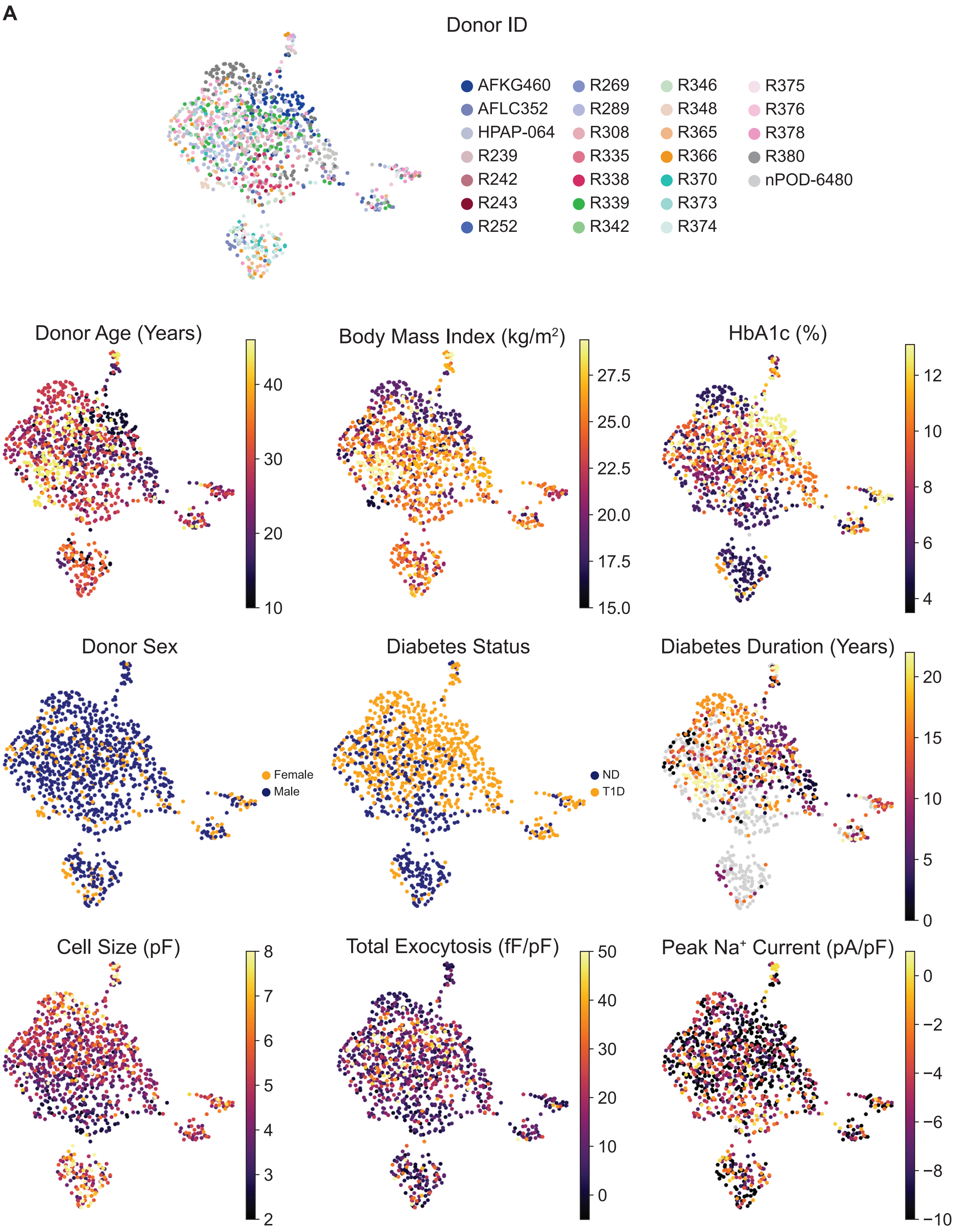

### Supplemental Figure 6

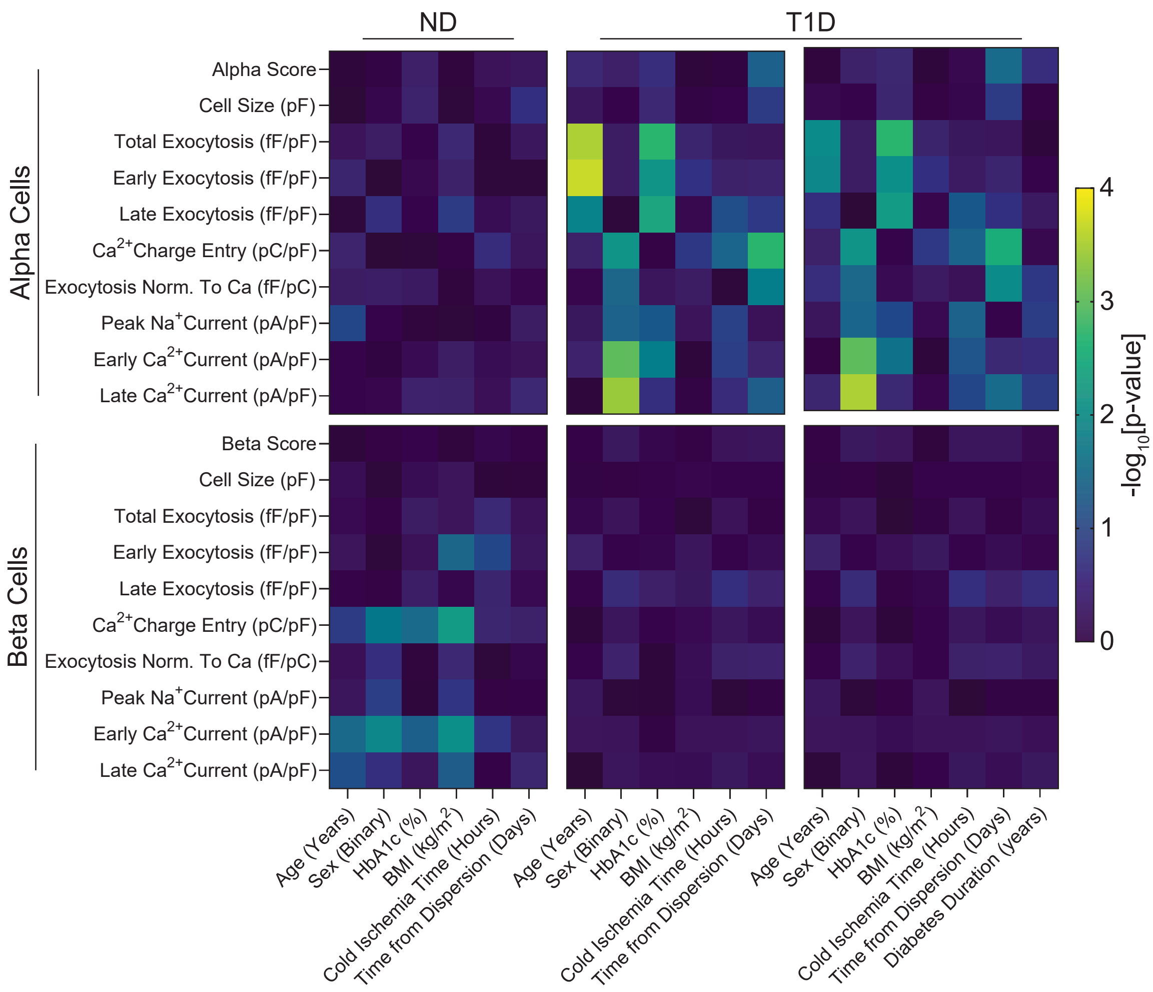

### Supplemental Figure 7

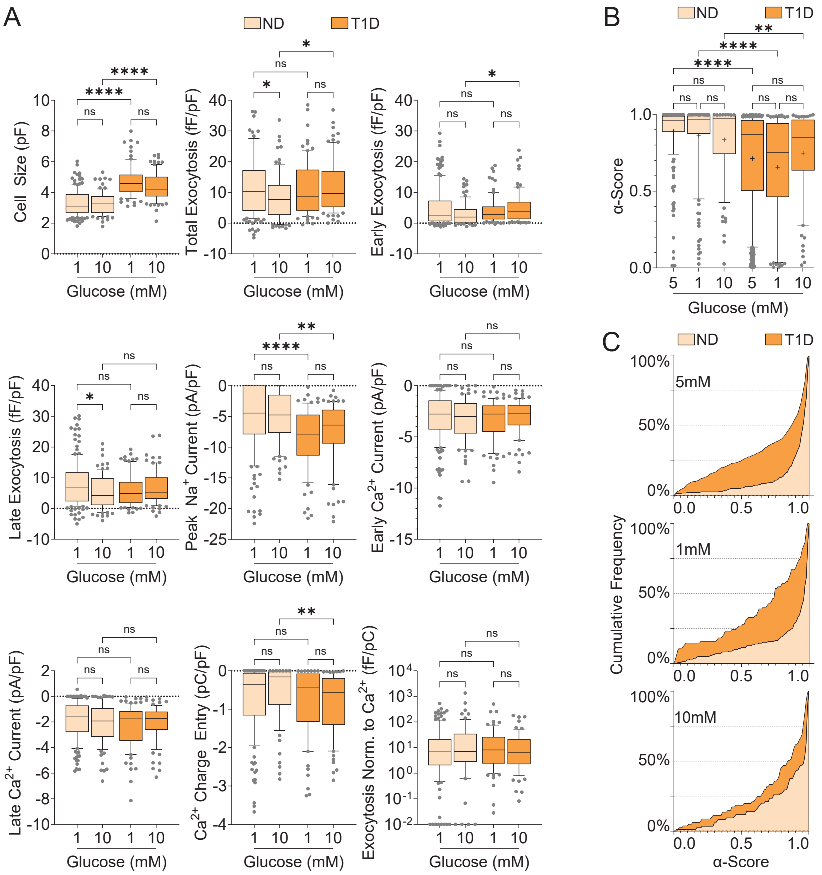

### Supplemental Figure 8

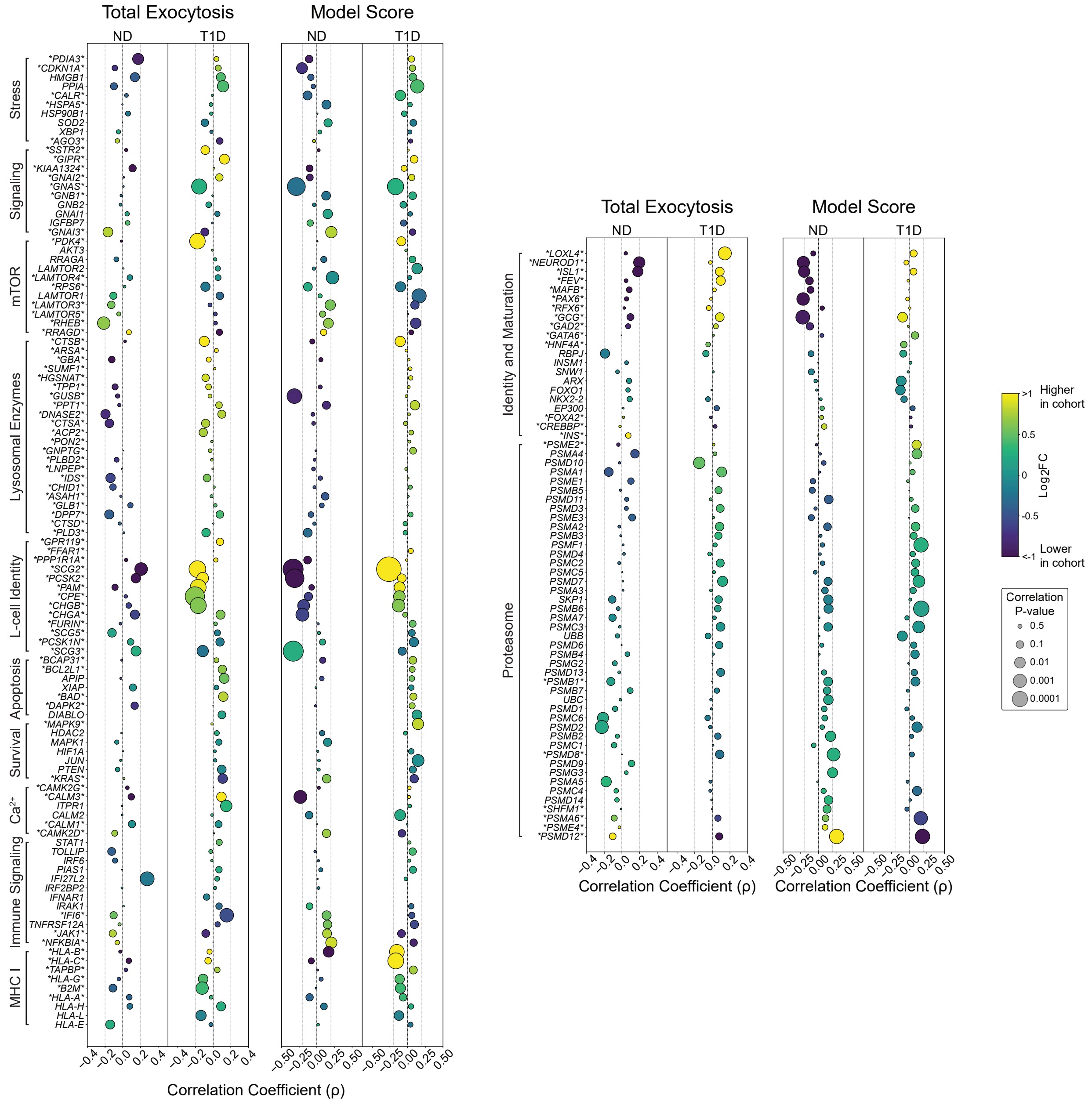

### Supplemental Figure 9

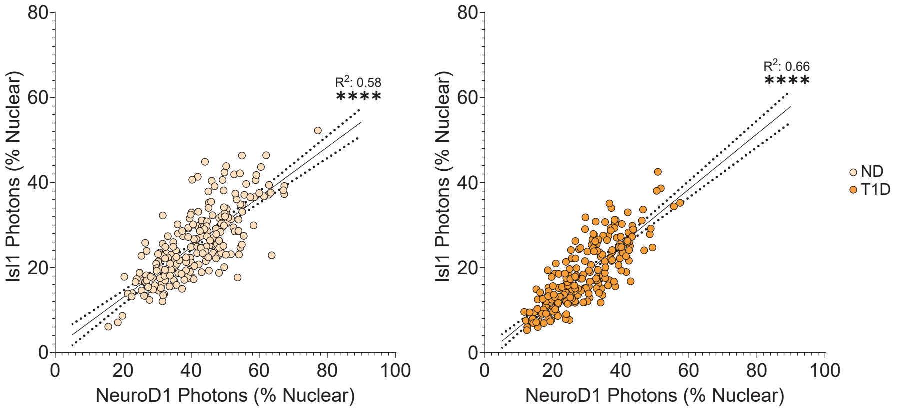
